# *Ex vivo* live imaging unveils the dynamics of oocyte formation in mice

**DOI:** 10.1101/2025.08.04.668457

**Authors:** Eishi Aizawa, Junko Hara, Takaya Abe, Tomoya S. Kitajima

## Abstract

Oocyte formation in mammals is a tightly regulated process essential for female fertility, yet the underlying mechanisms remain poorly understood. In this study, we establish an *ex vivo* culture system that faithfully recapitulates *in vivo* development and enables long-term live imaging of mouse fetal ovaries. Using high resolution imaging, we capture the dynamic behaviors of germ cells during the development from oogonia to nascent oocytes. We identify pronounced blebbing activity during the mitosis-to-meiosis transition. This behavior is regulated by meiotic initiation signals, underscoring its potential developmental relevance, although its precise role remains unclear. A prevailing model suggests that oocyte formation involves organelle transfer from neighboring germ cells during cyst breakdown. However, through photoconversion-based tracking, we observe no detectable transfer of mitochondria or centrosomes, as organelles remain confined to individual cells. These findings point to alternative mechanisms for cytoplasmic enrichment in oocytes. Our study provides new insights into mammalian oocyte formation and establishes a powerful platform for analyzing germ cell dynamics in real time.

## Introduction

Germ cells are fundamental to the continuity of life, ensuring the transmission of genetic information across generations. Among germ cells, oocytes play a vital role, as they contribute genetic material while also providing cytoplasmic components and organelles essential for early embryonic development. Despite their biological importance, the mechanisms underlying the formation of oocytes remain largely elusive. In mammals, germ cells originate as primordial germ cells (PGCs), which arise early in embryogenesis, around embryonic day (E) 7.25 (Saitou, 2021). These PGCs migrate to the developing gonads, where their fate is determined by local signaling cues. Around E11.5, bone morphogenetic protein (BMP) signaling in the gonadal environment directs these cells toward oogenic fate, prompting their differentiation into female-type germ cells known as oogonia (Miyauchi et al., 2017; Wu et al., 2016). Oogonia proliferate through mitotic divisions before entering meiosis, typically around E13.5-E14.5, and progressing to oocyte development (Spiller & Bowles, 2022).

A critical aspect of this developmental transition is the shift from mitotic proliferation to meiotic entry. This mitosis-to-meiosis transition is tightly regulated by extrinsic signals, notably the BMP and retinoic acid (RA) signaling (Bowles et al., 2006; Koubova et al., 2006; Miyauchi et al., 2017). BMP signaling confers meiotic competence to oogonia, while RA signaling induces the expression of *Stra8*, a transcription factor essential for initiating meiosis (Anderson et al., 2008). A recent study demonstrated that BMP signaling and STRA8 function cooperatively to ensure proper progression through the mitosis-to-meiosis transition (Cheung et al., 2025). In contrast to male germ cells, which enter a prolonged mitotic arrest, female germ cells proceed through a pre-meiotic S-phase and initiate meiosis during fetal development (Miles et al., 2010). However, the cellular dynamics accompanying the mitosis-to-meiosis transition remain poorly characterized, underscoring the need for further investigation.

Upon completing mitotic divisions, oogonia form germ cell cysts through incomplete cytokinesis, resulting in clusters of interconnected cells via intercellular bridges (Pepling & Spradling, 1998). A recent study showed that the balance between germ cell motility and the stability of intercellular bridges regulates the size of cysts (Levy et al., 2024). Within these cysts, a subset of germ cells undergoes meiotic progression, while others are eliminated during cyst breakdown, a process that separates interconnected germ cells into individual oocytes. Between E14.5 and postnatal day (PD) 5, approximately 80% of germ cells are lost through this process (Niu & Spradling, 2022). Following cyst breakdown, the surviving oocytes are individually enclosed by pregranulosa cells, forming primordial follicles beginning around PD1 (O’Connell & Pepling, 2021). These follicles then remain dormant for extended periods, over a year in mice and approximately 50 years in humans, ensuring a sustained ovarian reserve critical for lifelong fertility (Zhang et al., 2014).

The mechanisms underlying oocyte formation have been investigated across various species. In insects and lower vertebrates, precursor germ cells undergo synchronous mitotic divisions, giving rise to germline cysts interconnected by intercellular bridges (Büning, 1994; Matova & Cooley, 2001). In *Drosophila*, germline cysts are formed where 16 cyst cells are connected by incomplete cell divisions (Spradling et al., 2022). The intercellular bridges between these cyst cells facilitate the transfer of organelle-enriched cytoplasm from surrounding cells to the dominant oocyte, ensuring its development into the mature egg (Cox & Spradling, 2003; Mahowald & Strassheim, 1970). In mammals, organelle and cytoplasm transfer within cysts is believed to follow a similar pattern but with more complexity (Ikami et al., 2023; Lei & Spradling, 2013, 2016; Niu & Spradling, 2022). During cyst breakdown, a subset of germ cells undergo selection, ultimately contributing to the establishment of a finite ovarian reserve. The interaction between cyst germ cells during cyst breakdown is thought to involve intercellular bridges, facilitating the transfer of organelles and cytoplasm between cyst cells. Using a cell labeling strategy in mice, Lei and Spradling traced the cyst development and reported that organelle-enriched cytoplasm is transferred from nursing germ cells to dominant germ cells presumably via plasma membrane gaps between cyst germ cells (Lei & Spradling, 2013, 2016). This transfer has also been proposed to facilitate oocyte selection and contribute to the formation of an organelle-enriched structure, known as the Balbiani body. Single-cell RNA sequencing by Niu and Spradling has explored the molecular mechanisms regulating cytoplasmic exchange in these cysts, providing insights into the regulation of oocyte formation (Niu & Spradling, 2022). Collectively, these studies suggest that oocyte formation through cyst breakdown follows a conserved mechanism in mammals, akin to that observed in *Drosophila*.

However, while organelle and cytoplasm transfer are widely accepted as a key process in oocyte formation, studies of this transfer primarily employ static observations, which are limited in capturing the dynamic nature of this process (Ikami et al., 2023; Lei & Spradling, 2013, 2016). Additionally, the necessity of cyst structures in mammalian oocyte formation has been questioned by the *Tex14* knockout mouse model, in which females lacking intercellular bridges remain fertile (Greenbaum et al., 2009; Ikami et al., 2021). These findings suggest that organelle and cytoplasm transfer in the cyst may not be essential for oocyte formation. Furthermore, a recent preprint study proposed an alternative mechanism of oocyte formation, involving autophagy-driven oocyte competition, where dominant oocytes engulf the debris of sacrificed oocytes (Zhang et al., 2025). While this mechanism presents a new perspective on oocyte formation, further experimental evidence is needed to fully confirm its validity and understand the underlying mechanisms. In this study, we established an *ex vivo* culture system that enables live imaging of germ cell development in mouse gonads or ovaries, allowing real-time observation of oocyte formation. This novel application led to the discovery of unique blebbing behavior in germ cells during the mitosis-to-meiosis transition. Furthermore, we employed high-resolution 3D time-lapse tracking to investigate organelle transfer during oocyte formation. Unexpectedly, we found no detectable mitochondrial or centrosome transfer between germ cells, challenging the previous model of organelle transfer from nursing germ cells to dominant oocytes. Our findings highlight the utility of this *ex vivo* live imaging system for studying dynamic cellular processes and offer new insights into the mechanisms underlying oocyte formation.

## Results

### Establishment of an *Ex vivo* Culture System for Mouse Female Germ Cell Development

To investigate the development of mouse female germ cells, we established an *ex vivo* culture system that enables continuous monitoring of developmental processes (Figure 1A). Building upon a previously reported system for *in vitro* oocyte generation from pluripotent stem cells (Aizawa et al., 2023), we adapted it to culture fetal gonads or ovaries on coverslips within a vessel, facilitating long-term live imaging. For clarity, we refer to the developing reproductive organ as a ‘gonad’ until E13.5, before morphological sex differentiation is complete, and as an ‘ovary’ from E14.5 onward. The system was first applied to E12.5 gonads, a stage just prior to the mitosis-to-meiosis transition and oocyte formation. This *ex vivo* system allowed us to track gonadal cell development for up to 36 days (Figure 1B-C). In this study, cultured samples are labeled by their duration *ex vivo* (e.g., E12.5 + 7d refers to an E12.5 gonad cultured for 7 days). By day 21, distinguishable follicles emerged, with further expansion observed by day 35 (Figure 1B). Dissection of gonads cultured for 36 days confirmed the presence of germinal vesicles (GVs) in oocytes, a defining characteristic of growing and fully-grown oocytes (Figure 1C). These observations suggest the validity of our *ex vivo* culture system in monitoring female germ cell development and recapitulating the early stages of folliculogenesis.

**Figure 1.**
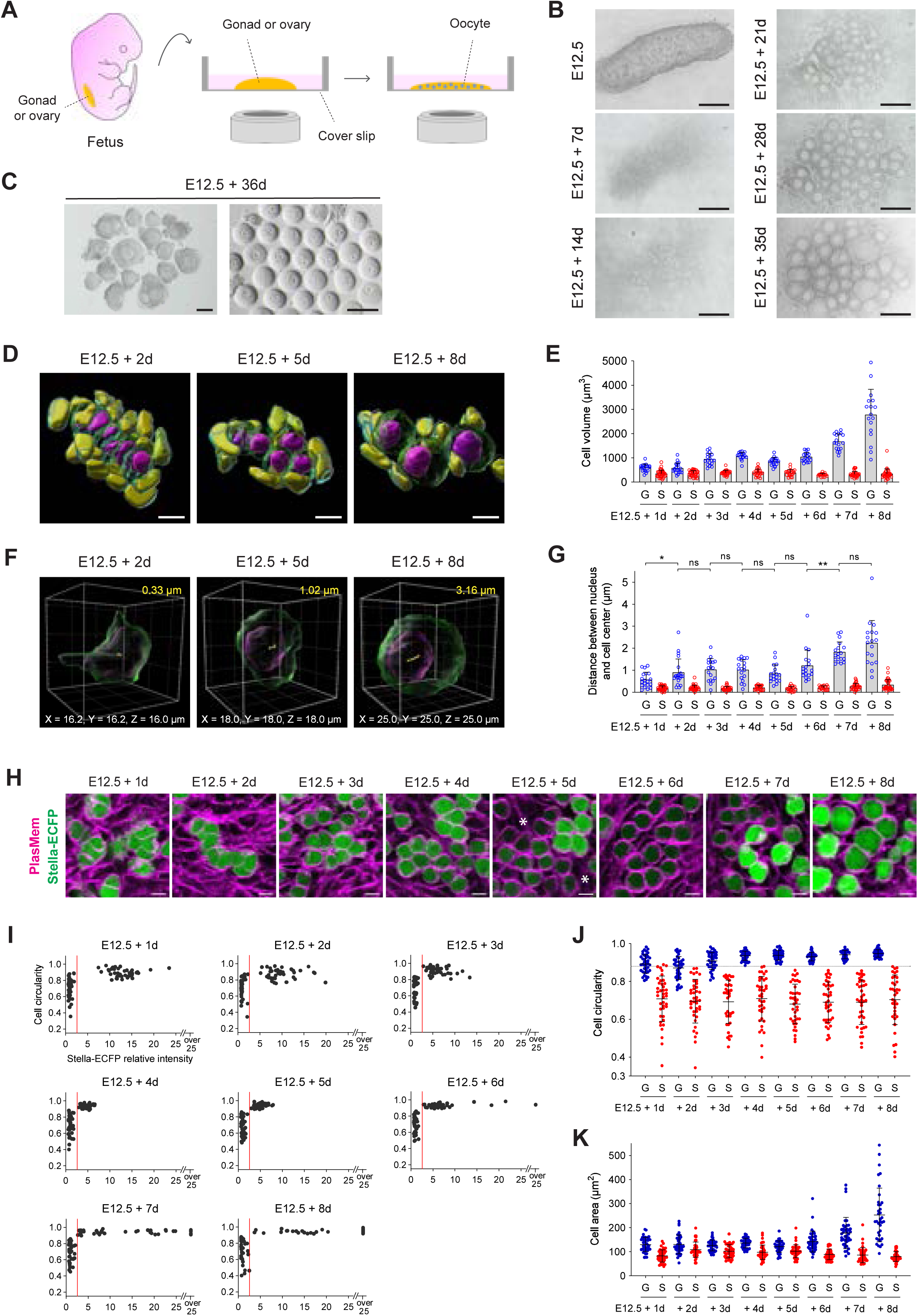
Establishment of an *ex vivo* culture system to visualize the development of mouse female germ cells. (A) A schematic illustration of the *ex vivo* culture system for visualizing the development of mouse female germ cells. A fetal gonad or ovary is cultured on a coverslip within the culture vessel, enabling microscopic observation of female germ cell development. Representative *ex vivo* development of an E12.5 gonad cultured within the vessel. Bright field images were captured at days 0, 7, 14, 21, 28 and 35. Follicles became distinguishable after 21 days of culture and further expanded by day 35. Scale bar, 200 μm. Bright field images of follicles (left) and oocytes (right) harvested from the E12.5 gonad shown in panel (B) and cultured *ex vivo* for 36 days. Germinal vesicles (GV) were present in each oocyte. Scale bar, 100 μm. (D) Representative three-dimensional reconstructed images of cells in E12.5 gonads expressing Stella-ECFP at 3, 5, and 8 days of *ex vivo* culture. Gonads were stained with Hoechst 33342 and PlasMem Bright Red. Germ cell membranes (green), germ cell nuclei (magenta), somatic cell membranes (cyan), and somatic cell nuclei (yellow) are shown. Confocal z-stacks were acquired at 1 μm intervals, and orthogonal 3D reconstructions were generated. Identical images are shown in Figure S1B. Scale bar, 10 μm. (E) Quantitative analysis of cell volume in cells from E12.5 gonads expressing Stella-ECFP during the 8-day *ex vivo* culture. Bars represent mean value + standard deviation. Volumes of germ cells (blue) and somatic cells (red) are plotted. Sample sizes (germ cells; somatic cells) were as follows: E12.5 + 1d (19; 36), E12.5 + 2d (21; 36), E12.5 + 3d (18; 26), E12.5 + 4d (18; 23), E12.5 + 5d (18; 21), E12.5 + 6d (18; 20), E12.5 + 7d (18; 31), and E12.5 + 8d (18; 34). G, germ cell; S, somatic cell. (F) Representative measurements of the distance between the nucleus and the cell surface center in germ cells from E12.5 gonads after 3, 5, and 8 days of *ex vivo* culture. Three-dimensional images were reconstructed from confocal z-stacks of gonads labeled with Stella-ECFP, Hoechst 33342, and PlasMem Bright Red. Germ cell membranes (green) and nuclei (magenta) are shown. Yellow spots indicate the centers of the cell membrane and the nucleus; the measured distance is shown in yellow. (G) Quantitative analysis of the distance between the nucleus and the cell surface center in cells from E12.5 gonads expressing Stella-ECFP during the 8-day *ex vivo* culture. Bars represent mean value + standard deviation. Distances for germ cells (blue) and somatic cells (red) are plotted. Sample sizes (germ cells; somatic cells) were as follows: E12.5 + 1d (19; 36), E12.5 + 2d (21; 36), E12.5 + 3d (18; 26), E12.5 + 4d (18; 23), E12.5 + 5d (18; 21), E12.5 + 6d (18; 20), E12.5 + 7d (18; 31), and E12.5 + 8d (18; 34). Statistical analysis was performed using a *t*-test with Welch’s correction. * *P* < 0.05; ** *P* < 0.01. G, germ cell; S, somatic cell. (H) Development of germ cells in E12.5 gonads expressing Stella-ECFP during an 8-day *ex vivo* culture. Gonads were stained with PlasMem Bright Red. The merged image shows Stella-ECFP expression (green) alongside the PlasMem signal (magenta). Stella-ECFP expression weakened around day 5 of culture but strengthened again by days 7 and 8. Asterisks indicate cells with weakened Stella-ECFP signals. Scale bar, 10 μm. (I) Quantitative analysis of Stella-ECFP intensity and cell circularity in cells observed from E12.5 gonads expressing Stella-ECFP during an 8-day *ex vivo* culture. Two distinct cell populations were identified: one with low Stella-ECFP intensity and low circularity, corresponding to somatic cells, and the other with high Stella-ECFP intensity and high circularity, representing germ cells. The mean Stella-ECFP intensity of the low population was normalized to a relative intensity of 1. Red lines indicate a Stella-ECFP relative intensity threshold of 2.5, used to distinguish germ cells (>2.5) from somatic cells (≤2.5). (J) Quantitative analysis of cell circularity in E12.5 gonads expressing Stella-ECFP during an 8-day *ex vivo* culture. Germ cells (blue; Stella-ECFP relative intensity >2.5) and somatic cells (red; Stella-ECFP relative intensity ≤2.5) were distinguished based on Stella-ECFP expression intensity. A circularity threshold of 0.88 (dashed line) also classified germ cells (>0.88) and somatic cells (≤0.88) between E12.5 + 4d and E12.5 + 8d. Bars represent mean values ± standard deviations. (K) Quantitative analysis of cell area in E12.5 gonads expressing Stella-ECFP during an 8-day *ex vivo* culture. Germ cells (blue; Stella-ECFP relative intensity >2.5) and somatic cells (red; Stella-ECFP relative intensity ≤2.5) were distinguished based on Stella-ECFP expression intensity. Bars represent mean values ± standard deviations. (I, J, K) Sample numbers for quantification (germ cells; somatic cells; gonads) were as follows: E12.5 + 1d (40; 40; 4), E12.5 + 2d (40; 40; 7), E12.5 + 3d (40; 40; 7), E12.5 + 4d (40; 40; 7), E12.5 + 5d (40; 40; 10), E12.5 + 6d (40; 40; 7), E12.5 + 7d (40; 40; 8), and E12.5 + 8d (40; 40; 8). G, germ cell; S, somatic cell.

### Characterization of Germ and Somatic Cell Morphology in *Ex vivo* Culture

We next characterized germ and somatic cell populations within the *ex vivo* culture system. To track their dynamics, we used *Stella*-ECFP reporter mice, in which the germline-specific gene *Stella* (also known as *Dppa3*) drives expression of a fusion protein consisting of STELLA and ECFP (Ohinata et al., 2008). The membrane-specific fluorescent dye, PlasMem Bright Red or Green, and Hoechst 33342 were also used to delineate individual cell boundaries and nuclei, respectively. Imaging of E12.5 gonads at daily intervals over an 8-day culture period captured morphological transitions in germ cells (Figure S1A). To comprehensively assess cell morphology during oocyte formation, we generated 3D images of cells each day throughout the 8-day culture period (Figure 1D, S1B). These 3D reconstructions captured the morphological changes of germ and somatic cells, illustrating the transition of germ cells from cyst-enclosed structures to individually spherical oocytes. Quantitative analysis of 3D morphology confirmed a rapid increase in germ cell volume after 6 days of culture, whereas somatic cell volume remained relatively unchanged (Figure 1E). We also measured the distance between the nuclear center and the cell surface center as a proxy for cell polarity (Figure 1F). While somatic cells maintained a consistently short distance, germ cells exhibited a significant increase, particularly after day 6 (Figure 1G). This result corroborates that germ cells acquire cell polarity following meiotic entry (Elkouby et al., 2016).

### Morphometric Strategy Enabling Marker-Independent Germ Cell Identification

To enhance the identification of germ and somatic cells in cultured tissues, we incorporated a morphometric analysis based on 2D images of E12.5 gonads cultured for 8 days (Figure 1H). Throughout this period, we observed a transient decrease in Stella-ECFP signal between days 5 and 7, followed by recovery on day 8, consistent with previous reports describing transient downregulation of *Stella* expression (Aizawa et al., 2023). Quantitative analysis revealed that Stella-ECFP intensity, when coupled with cell circularity, effectively distinguished the germ cell population from the somatic cell population (Figure 1I). A threshold Stella-ECFP relative intensity of 2.5 was sufficient to identify germ cells. Furthermore, cell circularity alone proved to be a reliable distinguishing factor, with a threshold of 0.88 effectively separating germ cells from somatic cells between days 4 and 8 of the culture (Figure 1J). In contrast, cell area was not a reliable parameter for distinguishing between germ and somatic cells (Figure 1K).

To further validate this approach, we analyzed nuclear morphology and histone intensity using Stella-ECFP and H2B-mCherry double-reporter mice (Figure S2A). Germ cells were again identified using a Stella-ECFP relative intensity threshold of 2.5. Quantitative assessment of nuclear characteristics demonstrated that H2B-mCherry intensity and nuclear circularity effectively distinguished these cell populations after 3 days of culture (Figure S2B). An H2B-mCherry intensity threshold of 0.63 identified germ cells (≤0.63) and somatic cells (>0.63) with over 98.8% accuracy between E12.5 + 3d and E12.5 + 6d (Figure S2C). Likewise, a nuclear circularity threshold of 0.93 reliably distinguished germ cells (≤0.93) from somatic cells (>0.93) with over 99.0% accuracy between E12.5 + 3d and E12.5 + 7d (Figure S2D). In contrast, consistent with cell area analysis (Figure 1K), nuclear area did not serve as a reliable parameter for distinguishing these cell populations (Figure S2E).

These analyses identified a set of morphological features that distinguish germ cells from somatic cells during oocyte formation. In addition to Stella-ECFP intensity, cell circularity, nuclear circularity, and histone intensity serve as reliable markers to distinguish germ cells from somatic cells beyond 3-days culture of E12.5 gonads. These markers were subsequently utilized to identify germ cells without reliance on Stella-ECFP reporters in the following experiments.

### Validation of Meiotic Progression in the *Ex vivo* Culture System

To validate our *ex vivo* culture system, we investigated meiotic progression by applying morphometric analysis using mCherry-SYCP3 (Enguita-Marruedo et al., 2018), a marker of chromosome axes, under both *in vivo* and *ex vivo* conditions. Germ cells were identified based on mCherry-SYCP3 signals up to E17.5 or E12.5 + 5d, and by a nuclear circularity exceeding 0.93 at later stages (Figure S2D). Imaging of an E12.5 + 6d gonad demonstrated chromosome axis formation comparable to those observed in an E18.5 ovary (Figure 2A). Additionally, E12.5 gonads cultured *ex vivo* ranging from 3 to 9 days were compared to ovaries from corresponding developmental stages from E15.5 to PD2 (Figure S3A-B). These comparisons demonstrated comparable patterns of chromosome axis formation, indicating that key meiotic processes are recapitulated in the *ex vivo* culture system. Quantitative analysis of chromosome axis formation ratios revealed no significant differences between *ex vivo* and *in vivo* samples at each corresponding time point, suggesting that meiotic progression occurs at a similar rate under both conditions (Figure 2B). These observations indicate that germ cells cultured *ex vivo* exhibit developmental characteristics comparable to their *in vivo* counterparts at equivalent time points (e.g., E12.5 + 3d cultured gonads resemble E15.5 ovaries). Together, these results support the use of the *ex vivo* culture system as a faithful model of female germ cell development.

**Figure 2.**
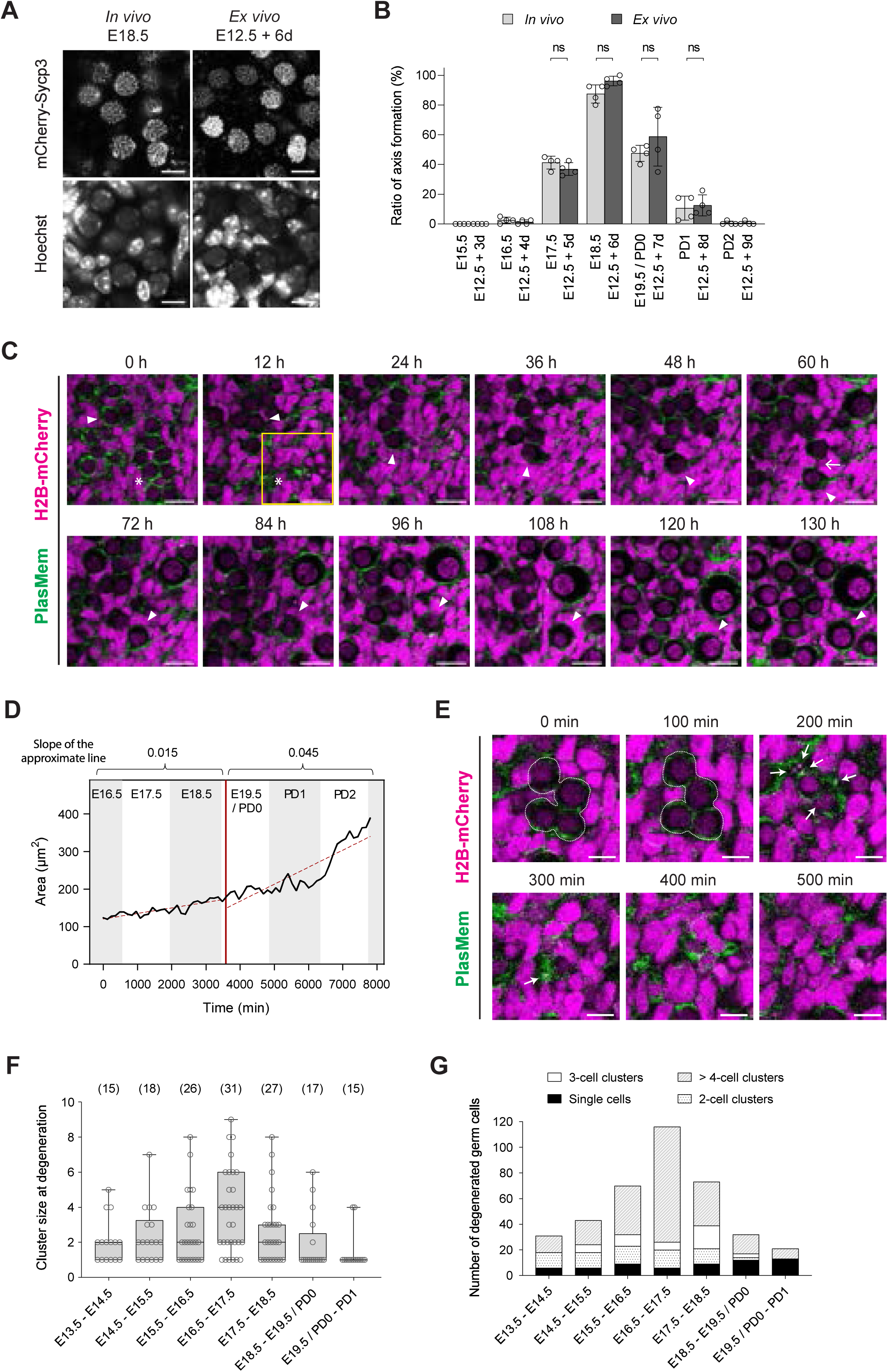
Application of the *ex vivo* culture system for live imaging analysis. (A) Representative images of cells in an E18.5 ovary and an E12.5 + 6d gonad, both expressing mCherry- SYCP3. Samples were stained with Hoechst 33342. Identical images are shown in Figure S3. Scale bar, 10 μm. (B) Quantitative analysis of chromosome axis formation ratios in germ cells from ovaries and E12.5 gonads cultured *ex vivo*, both expressing mCherry-SYCP3. Germ cells with or without distinct axis formation were evaluated based on mCherry-SYCP3 signals from confocal z-section images. *N* = 4 gonads/ovaries; 40 germ cells per gonad/ovary. Statistical analysis was performed using a *t*-test with Welch’s correction. ns, non-significant. (C) Representative live imaging of oocyte formation. An E12.5 + 4d gonad expressing H2B-mCherry was stained with PlasMem Bright Green, and imaged every 10 minutes continuously for 130 hours. PlasMem (green) and H2B-mCherry (magenta) signals are shown as merged images. Arrowheads mark tracked germ cells/oocytes. Asterisks indicate a degenerated cluster of germ cells, and an arrow marks the separation of the germ cell from a cluster. The region enclosed by the yellow square is shown in Figure 2E at higher magnification and temporal resolution. Scale bar, 20 μm. See also Movie 1. (D) Time-lapse quantification of the area of the germ cell tracked in (C). Cell area was measured every 120 minutes over a 130-hour period. The red line marks the time point at which the germ cell separated from the cluster. Dotted lines indicate approximate trends in cell area before and after separation. The slope of each trend line is shown above the plot. *Ex vivo* time points are converted to corresponding *in vivo* developmental times, as shown at the top of the plot. (E) Time-lapse view of the region marked in (C), showing synchronized degeneration of a germ cell cluster at higher magnification and finer temporal resolution. Images were extracted from the same time-lapse dataset as in (C) and are shown at 100-minute intervals. Dotted lines outline the germ cell cluster; arrows indicate particles remaining after degeneration. See also Movie 1. (F) Quantification of germ cell cluster size at the time of degeneration in E12.5 gonads cultured *ex vivo* for 1 to 8 days. Degenerating germ cells were counted for each event within 12 randomly selected 100 μm × 100 μm regions per developmental window. The data are plotted according to the corresponding *in vivo* developmental ages (E13.5 to PD1), converted from *ex vivo* time points (E12.5 + 1d to E12.5 + 8d). Data points (circles) represent the cluster sizes observed at individual degeneration events. Box and whisker plots show the 25th, 50th (median), and 75th percentiles (box), with whiskers indicating the minimum and maximum values. Numbers in brackets indicate the number of degeneration events observed at each developmental window. For each developmental window, six E12.5 gonads cultured *ex vivo* were analyzed for quantification. (G) Cumulative number of degenerated germ cells in E12.5 gonads cultured *ex vivo* for 1 to 8 days, categorized by cluster size at the time of degeneration. Degeneration events were grouped into four categories based on the number of germ cells per cluster: single cells, 2-cell clusters, 3-cell clusters, and clusters of more than 4 cells. Developmental ages correspond to the *in vivo* timeline (E13.5 to PD1), converted from *ex vivo* time points (E12.5 + 1d to E12.5 + 8d). Quantification was based on the same dataset used in (F). For each developmental window, six E12.5 gonads cultured *ex vivo* were analyzed.

### Autonomous Expansion and Synchronous Degeneration of Germ Cells

To comprehensively capture oocyte formation dynamics, we performed long-term live imaging of E12.5 + 4d gonads cultured *ex vivo*. At this stage, germ cells were identified by their spherical morphology and low H2B-mCherry fluorescence intensity (Figure 1J, S2C). Continuous imaging of gonads expressing H2B- mCherry and stained with PlasMem Bright Green allowed tracking of individual germ and somatic cells throughout oocyte formation over a 5-day period (Figure 2C, Movie 1).

We observed a marked increase in germ cell size over time, with cell area expanding more than threefold during the imaging period (Figure 2C-D). Notably, this expansion occurred autonomously after cells separated from the cluster—presumably a cyst. The rate of area expansion increased approximately threefold following separation compared to the period prior. This observation contrasts with the prevailing model that oocytes increase in size through cytoplasmic transfer within the germ cell cyst (Lei & Spradling, 2016). Instead, our findings align with a recent preprint suggesting that cytoplasmic enrichment occurs through the engulfment of debris from degenerated oocytes (Zhang et al., 2025).

In addition to cell growth, we frequently observed the synchronous degeneration of germ cells within individual clusters (Figure 2C, E), often marked by residual histone signals indicative of apoptosis (Lei & Spradling, 2013). To quantify this process, we counted degenerating germ cells in 12 randomly selected 100 μm × 100 μm regions from E12.5 gonads cultured *ex vivo*, corresponding to *in vivo* stages E13.5 to PD1 (Figure 2F). Degeneration of multicellular clusters was commonly observed throughout this period. Cluster size peaked between E16.5 and E17.5, with a median of four germ cells per degenerating cluster.

We further categorized degeneration events by cluster size and quantified the cumulative number of degenerated germ cells (Figure 2G). Across the observed time window, single cell degeneration accounted for only 15.8% of all degeneration events. In contrast, multicellular degeneration—clusters containing three or more germ cells—represented the majority of events particularly during E15.5–E16.5 (67.1%), E16.5– E17.5 (82.8%), and E17.5–E18.5 (71.2%). While previous studies have described a gradual reduction in germ cell number within cysts via selective nurse cell death (Ikami et al., 2023; Lei & Spradling, 2016; Niu & Spradling, 2022), our live imaging revealed that germ cells within a cluster—likely corresponding to germ cell cysts—appear to be eliminated simultaneously. Our findings highlight synchronous cluster degeneration as an underrecognized mode of cyst breakdown during oocyte formation (see Discussion).

### Quantitative Analysis of Germ Cell Nuclear Rotation

To further characterize subcellular behaviors during oocyte formation, we analyzed the dynamics of nuclear movement in germ cells. Nuclear rotation has been demonstrated to be involved in chromosome movement in spermatocytes (Lee et al., 2015; Shibuya et al., 2014) and is proposed to play a role in the maintenance of oocyte dormancy in mice (Nagamatsu et al., 2019). Time-lapse imaging at 10-second intervals captured the rotational movements of germ cell nuclei in E12.5 + 4d gonads stained with Hoechst 33342 (Figure S4A, Movie 2). Quantification of nuclear rotation speed across multiple germ cells revealed considerable cell-to-cell variability, with some nuclei exhibiting rapid movements, while others moved more gradually (Figure S4B-D). Velocity plots further illustrated the directionality of nuclear rotation, with data distributed across all quadrants, ranging from 12.9% to 35.5% (Figure S4E-G). These findings indicate that nuclear rotation occurs in multidirectional rather than restricted to a defined axis.

### Dynamic Blebbing in Germ Cells During the Mitosis-to-Meiosis Transition

The mitosis-to-meiosis transition represents a pivotal developmental shift in germ cell development; however, the extent to which this process involves specific changes in cellular behavior remains poorly understood. To investigate potential morphological features associated with this transition, we performed live imaging with high temporal resolution during the critical window encompassing the mitosis-to-meiosis shift. Unexpectedly, we frequently observed dynamic blebbing activity in germ cells from E12.5 + 2d gonads, characterized by transient protrusions forming and retracting at various regions of the cell surface (Figure 3A-B, Movie 3). To quantify this behavior, we measured the number of blebs with a height exceeding 2 µm from E11.5 to E12.5 + 5d (Figure 3C). Only female gonads were used for this analysis, with sex at E11.5 determined by genotyping (Figure S5A). The analysis revealed that blebbing frequency increased as development progressed, peaking at E12.5 + 2d at a rate exceeding one bleb per cell per minute, followed by a marked decline by E12.5 + 3d (Figure 3D). This temporal pattern coincided with the mitosis-to-meiosis transition, which occurs around E13.5-E14.5 (Ishiguro, 2023), suggesting a potential link between blebbing activity and early meiotic events. Notably, blebbing was exclusively observed in the direction from germ cells to adjacent somatic cells, with rare instances of blebbing between germ cells (Figure 3E). This directional bias implies differential regulation of membrane dynamics at homotypic versus heterotypic interfaces, thereby restricting blebbing to the latter. Since blebbing is often linked to fundamental cellular processes such as apoptosis, cell migration, and cytokinesis, its occurrence in germ cells may reflect an active developmental transition (Ikenouchi & Aoki, 2022). Given that a subset of germ cells remains mitotically active at this stage, we initially hypothesized that blebbing might be associated with cytokinetic activity, as previously reported (Burton & Taylor, 1997; Wang et al., 2021). However, contrary to this assumption, blebbing was absent in germ cells undergoing cytokinesis, whereas neighboring germ cells displayed active blebbing in the absence of cytokinetic activity (Figure S5B, Movie 4).

**Figure 3.**
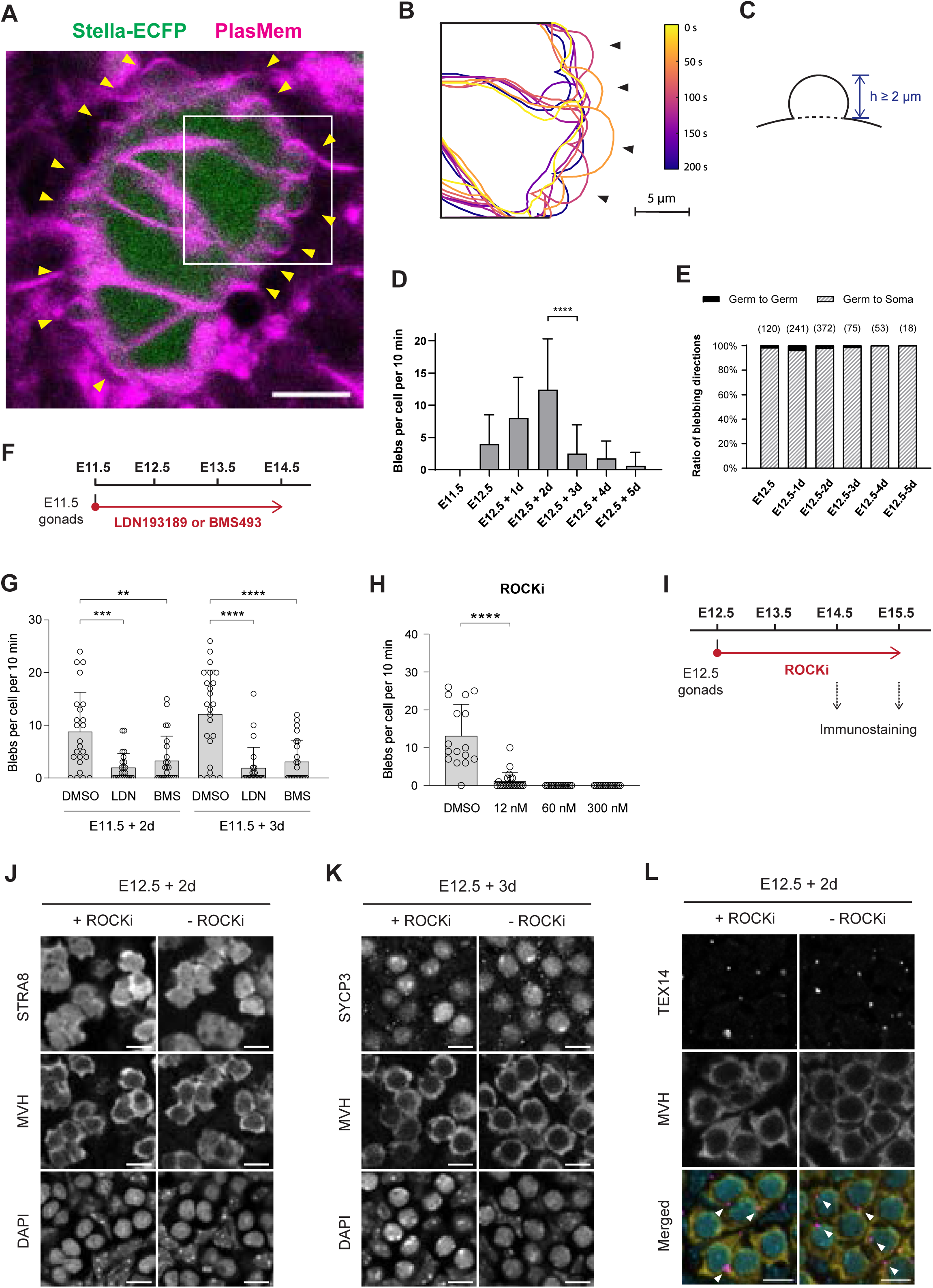
Dynamics of blebbing in germ cells observed during the mitosis-to-meiosis transition. (A) Representative image of cells in an E12.5 gonad expressing Stella-ECFP after 2 day of *ex vivo* culture. The gonad was stained with PlasMem Bright Red. Stella-ECFP (green) and PlasMem (magenta) signals are shown as a merged image. Arrowheads indicate blebs on germ cells. The time-course projection within the white square is shown in (B). Scale bar, 20 μm. See also Movie 3. (B) Representative projection of germ cell membranes in an E12.5 gonad expressing Stella-ECFP after 2 days of *ex vivo* culture. Germ cell membranes stained with PlasMem Bright Red were traced from live imaging captured every 2 seconds. The projection shows seven tracings over 200 seconds. Arrowheads indicate blebs. See also Movie 3. (C) Schematic illustration of a bleb. A bleb is defined as an expanded region of the cell membrane with a height (*h*) exceeding 2 μm. (D) Blebbing frequency over the developmental timeline. Blebs with heights exceeding 2 μm were counted for each germ cell in female gonads expressing Stella-ECFP and stained with PlasMem Bright Red. The sex of E11.5 embryos was determined by PCR genotyping (Figure S5A), and only female gonads were analyzed. *N* = 30 germ cells per developmental time point. Bars represent mean values + standard deviations. Statistical analysis was performed using a *t*-test with Welch’s correction. **** *P* < 0.0001. (E) Direction of blebbing over the developmental timeline. Blebbing was categorized as either germ cell- to-germ cell or germ cell-to-somatic cell. No blebbing from somatic cells was observed. Numbers in brackets indicate the total number of blebbing observed at each developmental time point. *N* = 30 germ cells per developmental time point. (F) Schematic timeline of the *ex vivo* culture to evaluate the effects of LDN1931189 or BMS493 on blebbing. Female E11.5 gonads were cultured with either inhibitor for 3 consecutive days. (G) Blebbing frequency following treatment with LDN193189 (500 nM) or BMS493 (10 μM). Bars represent mean values + standard deviations. *N* = 24 germ cells per condition. Statistical analysis was performed using a *t*-test with Welch’s correction. ** *P* < 0.01; *** *P* < 0.001; **** *P* < 0.0001. See also Figure S6A and Movie 6. (H) Blebbing frequency following treatment with ROCKi for 16 hours. The number of blebs was counted from live imaging of E12.5 +2d gonads expressing Stella-ECFP. Bars represent mean values + standard deviations. Statistical analysis was performed using a *t*-test with Welch’s correction. Sample sizes: DMSO (*n* = 16), 12 nM (*n* = 23), 60 nM (*n* = 16), 300 nM (*n* = 16). **** *P* < 0.0001. (I) Schematic timeline of the *ex vivo* culture to evaluate the effect of ROCKi on meiosis initiation. E12.5 gonads were cultured in the presence of ROCKi (12 nM) for 2 or 3 consecutive days, followed by immunostaining analysis. (J, K) Representative immunostaining of meiosis markers in cultured gonads following ROCKi treatment. (J) E12.5 + 2d gonads stained with STRA8 and MVH antibodies. (K) E12.5 + 3d gonads stained with SYCP3 and MVH antibodies. Both STRA8 and SYCP3 signals were detected in germ cells irrespective of ROCKi treatment. Scale bar, 10 μm. (L) Representative immunostaining of TEX14 in cultured gonads following ROCKi treatment. E12.5 + 2d gonads were stained with antibodies against TEX14 and MVH, followed by DAPI counterstaining. Merged images show TEX14 (magenta), MVH (yellow) and DAPI (cyan). TEX14 signals (arrowheads) were detected between germ cells irrespective of ROCKi treatment. Scale bar, 10 μm.

To investigate the mechanisms underlying blebbing in germ cells, we analyzed cytoskeletal dynamics (Figure S5C, Movie 5). Co-staining of actin filaments with SiR-Actin and a membrane marker revealed that actin accumulation was associated with bleb contraction. To quantify blebbing dynamics, we measured the bleb arc length, which increased during expansion and shortened during contraction (Figure S5D-E). Analysis of fluorescence intensity along the bleb arc showed that PlasMem Bright Red intensity remained stable throughout the blebbing process, whereas SiR-Actin intensity increased as blebbing transitioned into the contraction phase. (Figure S5F). These results indicate that actin dynamics contribute to bleb retraction in germ cells.

### Regulation of Blebbing by Meiotic Initiation Signals and Cytoskeletal Dynamics

To elucidate the molecular mechanisms underlying blebbing, we investigated the effects of specific signaling inhibitors. Given that blebbing activity peaked during the mitosis-to-meiosis transition (Figure 3D), we first examined its potential correlation with meiotic initiation pathways, specifically BMP and RA signaling (Ishiguro, 2023; Spiller & Bowles, 2022). Previous studies have demonstrated that *ex vivo* culture of E11.5 gonads with LDN193189, an ALK2/3 receptor inhibitor, or BMS493, an RA receptor antagonist, disrupts meiotic initiation (Miyauchi et al., 2017). Following a similar approach (Figure 3F), we observed that treatment with these inhibitors significantly reduced blebbing frequency in both E11.5 + 2d and E11.5 + 3d gonads (Figure 3G, S6A, Movie 6). These results indicate that blebbing is regulated by meiotic initiation pathways through BMP and RA signaling.

We next investigated the role of cytoskeletal dynamics, prompted by observations of actin involvement (Figure S5C-F). Treatment with Jasplakinolide (an actin polymerization stabilizer; IC_50_ ≈ 15-170 nM) (Bubb et al., 1994; Senderowicz et al., 1995), Cytochalasin D (an actin polymerization inhibitor; IC_50_ ≈ 25-150 nM) (Luxenburg et al., 2012; Sayyad et al., 2015), and Ciliobrevin D (a dynein inhibitor; IC_50_ ≈ 15-50 µM) (Firestone et al., 2012; Tati & Alisaraie, 2021) did not suppress blebbing at concentrations within the IC_50_ range reported in other cell types (Figure S6B-D). Although higher doses of Jasplakinolide (1 µM) and Cytochalasin D (500 nM) reduced blebbing, these conditions also induced significant cell death, suggesting that the observed suppression was possibly due to cytotoxic effects rather than direct inhibition of blebbing (Figure S6E). To further dissect the underlying mechanisms, we evaluated the roles of actin polymerization and contractility. Inhibition of the Arp2/3 complex (CK666) and formin-mediated actin polymerization (SMIFH2) had minimal impact on blebbing (Figure S6F-G). In contrast, inhibition of myosin II using Blebbistatin markedly suppressed blebbing (Figure S6H), indicating that actomyosin contractility is essential for this process, consistent with findings in other cell types (Ikenouchi & Aoki, 2022). Given the role of myosin II, we further examined Rho-associated coiled-coil kinase (ROCK), which promotes actin polymerization via mDia/formin activation and enhances myosin II activity by inhibiting myosin light chain phosphatase (Amano et al., 2010). Treatment with the ROCK inhibitor H1152 significantly reduced blebbing frequency in E12.5 + 2d gonads without affecting cell viability (Figure 3H, S6I). These findings underscore the importance of cytoskeletal regulation, particularly actomyosin contractility and ROCK signaling, in germ cell blebbing (Figure 3H).

Finally, we assessed whether blebbing is functionally linked to meiotic progression. Notably, despite continuous ROCK inhibition from E12.5 onward (Figure 3I), meiotic markers STRA8 (E12.5 + 2d) and SYCP3 (E12.5 + 3d) remained expressed (Figure 3J-K), indicating that meiosis proceeds independently of blebbing. Similarly, TEX14 signals, associated with interconnected cyst formation, remained unaffected by ROCK inhibition (Figure 3L). These findings suggest that while blebbing is regulated by RA and BMP signaling, it represents a process distinct from meiotic initiation. Given that blebbing is involved in various fundamental cellular processes, its occurrence in germ cells may reflect an active developmental transition. However, the precise functional significance of blebbing in oocyte formation remains to be elucidated.

### Absence of Detectable Mitochondrial Transfer Revealed by Live Imaging

Organelle transfer between germ cells has been proposed as a key mechanism in oocyte formation, enabling the enrichment of cellular components essential for oocyte development. Lei and Spradling suggested that mitochondria, Golgi complexes, centrosomes, and other cytoplasmic materials are transferred from sister cyst germ cells to the developing oocyte, primarily between E16.5 and E19.5/PD0 (Lei & Spradling, 2016). This intercellular transfer has also been implicated in the assembly of the Balbiani body, an organelle-dense structure characteristic of early oocytes. However, direct evidence supporting such transfer remains limited. To better understand this mechanism, we utilized our live imaging system to investigate the dynamics of organelle transfer during oocyte formation.

We first performed live imaging of Golgi complexes using Golgi-EGFP, in which EGFP is fused to the N-terminal region of β-1,4-galactosyltransferase 1 (Figure S7A, Movie 7) (Abe et al., 2011). This reporter effectively labeled Golgi complexes, and we successfully tracked the formation of the Golgi ring, a hallmark of the Balbiani body. However, the ubiquitous expression of the reporter across all cells made it difficult to discern potential intercellular transfer events. We then imaged mitochondrial dynamics using Mito-EGFP, which labels mitochondria via fusion to the N-terminal region of cytochrome c oxidase subunit VIII A (Figure S7B, Movie 8) (Abe et al., 2011). Although mitochondrial movements and germ cell disruption were clearly captured, evidence of intercellular transfer remained inconclusive due to the ubiquitous expression of the reporter, similar to what was observed with Golgi-EGFP.

To enhance detection, we employed Mito-Dendra2 mice, which express Dendra2, a photoconvertible reporter (Pham et al., 2012). Mitochondria in a targeted germ cell were photoconverted from green to red fluorescence, followed by 3D time-lapse tracking and imaging to capture potential transfer to adjacent germ cells (Figure 4A-B). As expected, the distinct red fluorescence facilitated precise mitochondrial tracking (Figure 4C, Movie 9). Despite the high resolution and improved visualization of our time-lapse imaging, we did not observe detectable mitochondrial transfer between germ cells across various time points, a total of 345 hours, including observation of 15 photoconverted germ cells and 24 adjacent, non-photoconverted germ cells. Quantification of photoconverted Red Mito-Dendra2 intensity further confirmed the absence of detectable mitochondrial transfer (Figure 4D-E; Figure S8). In our comprehensive time-lapse analysis, we tracked the mean intensity of photoconverted Red Mito-Dendra2 in both the photoconverted germ cells and adjacent, non-photoconverted germ cells. Throughout the imaging period, no significant increase in Red Mito-Dendra2 intensity was observed in adjacent germ cells, indicating that mitochondrial transfer is not a prominent event during this stage of oocyte formation.

**Figure 4.**
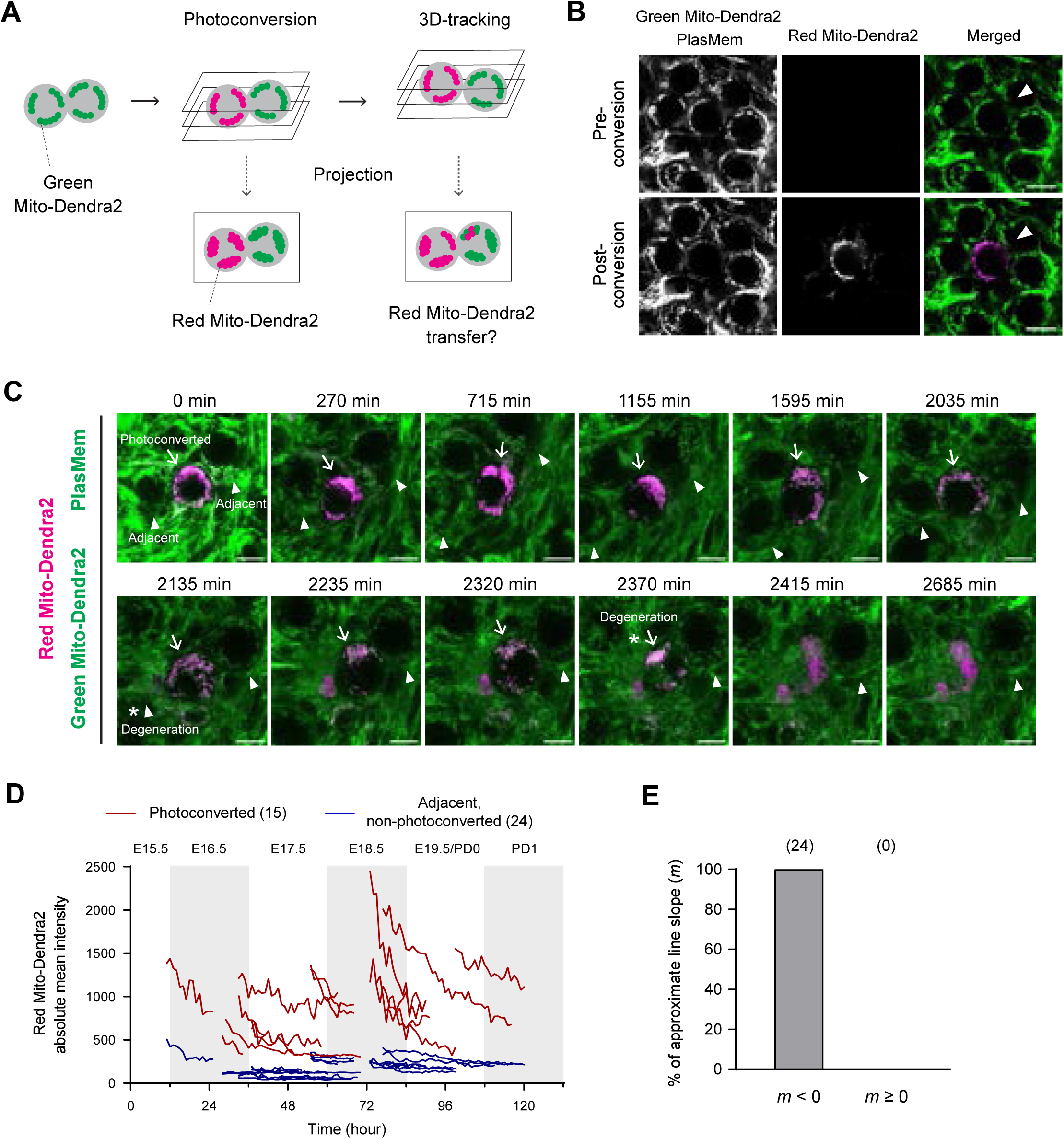
Non-detectable mitochondrial transfer between germ cells during oocyte formation. (A) Schematic strategy for assessing mitochondrial transfer between germ cells. Green Mito-Dendra2 is photoconverted to Red Mito-Dendra2 in a targeted germ cell, followed by 3D time-lapse imaging of both the targeted and adjacent germ cells. The targeted cell is tracked during imaging based on the Red Mito-Dendra2 intensity. Captured 3D images are projected onto 2D planes to evaluate Red Mito- Dendra2 signals in adjacent germ cells. Representative images of Mito-Dendra2 photoconversion. Green Mito-Dendra2 in a targeted germ cell (arrowheads) was photoconverted to Red Mito-Dendra2 in an E12.5 + 6d gonad expressing Mito-Dendra2, following staining with PlasMem Bright Green. Scale bar, 10 μm (C) Representative live imaging of an E12.5 + 6d gonad expressing Mito-Dendra2. The gonad was stained with PlasMem Bright Green and imaged at 5-minute intervals. Merged images show PlasMem (green), non-photoconverted Green Mito-Dendra2, and photoconverted Red Mito-Dendra2 signals. Arrows indicate a photoconverted germ cell, and arrowheads mark adjacent non-photoconverted germ cells. Asterisks denote the onset of germ cell degeneration. Scale bar, 10 μm. See also Movie 9. (D) Comprehensive time-lapse quantification of photoconverted Red Mito-Dendra2 intensity during oocyte formation. Green Mito-Dendra2 in a germ cell within cultured gonads/ovaries was photoconverted to Red Mito-Dendra2, followed by 3D tracking and imaging at 5-minute intervals. Absolute mean intensities of Red Mito-Dendra2 in photoconverted germ cells and adjacent non-photoconverted germ cells were measured and aligned by developmental time. *Ex vivo* time points were converted to corresponding *in vivo* developmental times, as shown at the top of the plot. *N* = 15 photoconverted germ cells and 24 adjacent non-photoconverted germ cells. See also Figure S8. (E) Summary of changes in Red Mito-Dendra2 intensity. Approximate lines on the plot of time versus Red Mito-Dendra2 intensity (D) were calculated for respective non-photoconverted germ cells adjacent to photoconverted germ cells. The slopes of these lines, denoted as *m*, were categorized into two groups: *m* < 0 or *m* ≥ 0. *N* = 24 slopes. Numbers in brackets indicate the sample sizes for each category.

### Characterization of Centrosomes during Oocyte Formation

Given that mitochondrial transfer was not detected, we further examined centrosomes, which are visualized as discrete foci rather than clusters, allowing for more precise tracking of potential organelle transfer between germ cells. Centrosomes, marked by centrin 2 (CETN2), were analyzed using EGFP-CETN2 mice, which express a fusion protein of human CETN2 tagged with EGFP, driven by the chicken beta-actin promoter (Higginbotham et al., 2004). This approach enabled us to assess centrosome dynamics at a single-organelle resolution during oocyte formation. To characterize centrosome distribution, we first quantified the number of EGFP-CETN2 foci at different developmental stages. Imaging of E12.5 gonads cultured *ex vivo* revealed considerable variation in centrosome numbers, which we categorized into single pair/cluster and two or more pairs/clusters (Figure S9A). Quantitative analysis showed a rapid increase in the proportion of germ cells exhibiting two or more pairs/clusters from E17.5 to E19.5/PD0 (Figure 5A), consistent with reports of pericentrin and γ-tubulin volume expansion during this period (Lei & Spradling, 2016). Centrosome distribution relative to germ cell diameter did not exhibit rapid changes, suggesting that developmental timing plays a more significant role in centrosome number (Figure S9B). Time-lapse 3D imaging allowed precise visualization of EGFP-CETN2 foci within individual germ cells (Figure S9C, Movie 10). However, similar to Golgi complex and mitochondrial tracking, the widespread presence of EGFP- CETN2 signals across cells precluded clear identification of centrosome transfer between adjacent cells in the absence of targeted labeling. To address this, we assessed the stability of EGFP-CETN2 signals using fluorescence recovery after photobleaching (FRAP) (Figure S9D-E). Photobleached EGFP-CETN2 signals did not recover significantly within 2 hours, indicating low turnover of CETN2 fluorescence and justifying a photoconversion-based approach. Thus, we employed a strategy akin to Mito-Dendra2 tracking, using a photoconvertible centrosome marker to directly monitor potential intercellular transfer (Figure 5B).

**Figure 5.**
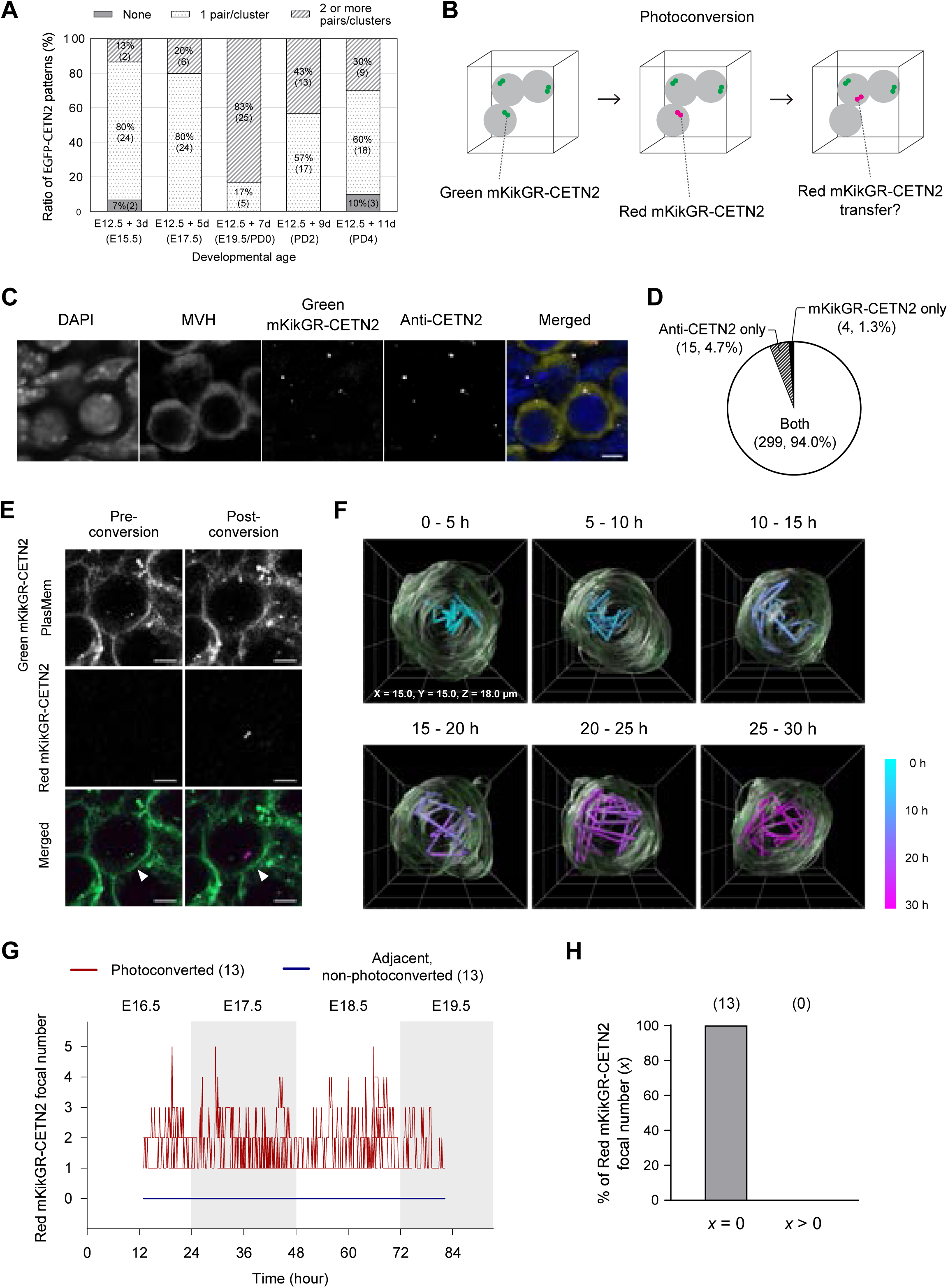
Generation of mKikGR-CETN2 mice and non-detectable centrin transfer between germ cells. (A) Distribution of EGFP-CETN2 patterns in germ cells across developmental age. EGFP-CETN2 patterns in each germ cell of E12.5 gonads cultured for 3 to 11 days were classified into three categories: None, 1 pair/cluster, and 2 or more pairs/clusters. The X-axis shows the corresponding *in vivo* developmental ages in brackets, inferred from *ex vivo* culture durations. Numbers in brackets within the bars indicate germ cell counts. *N* = 30 germ cells per developmental age. See also Figure S9A-B. (B) Schematic of the strategy for assessing centrin transfer between germ cells. Green mKikGR-CETN2 is photoconverted to Red mKikGR-CETN2 in a targeted germ cell, followed by 3D time-lapse imaging of both the targeted and adjacent germ cells. Centrin transfer is indicated by the appearance of Red mKikGR-CETN2 in an adjacent germ cell. (C) Representative immunostaining of a mKikGR-CETN2 ovary. E17.5 ovaries expressing mKikGR-CETN2 were stained with MVH and CETN2 antibodies and counterstained with DAPI. The merged image shows DAPI (blue), MVH (yellow), mKikGR-CETN2 (green) and anti-CETN2 (magenta). Scale bar, 5 μm. (D) Summary of centrin immunostaining in mKikGR-CETN2mouse ovaries. Green mKikGR-CETN2 and anti- CETN2 foci were assessed in E17.5 ovaries expressing mKikGR-CETN2. Focal patterns were classified into three categories: Both (co-localization of mKikGR-CETN2 and anti-CETN2), mKikGR-CETN2 only, and
anti-CETN2 only. Numbers in brackets indicate foci counts and their proportions among all foci. Centrin signals were analyzed in a total of 226 germ cells from 3 ovaries. (E) Representative images of mKikGR-CETN2 photoconversion. An E12.5 + 5d gonad expressing mKikGR-CETN2 was stained with PlasMem Bright Green, followed by 405 nm exposure targeting a centrin signal (arrowheads). In the merged image, PlasMem and unconverted Green mKikGR-CETN2 are shown in green, while photoconverted Red mKikGR-CETN2 is shown in magenta. Scale bar, 5 μm. (F) Three-dimensional reconstructed images of a germ cell with photoconverted Red mKikGR-CETN2 signals. An E12.5 + 5d gonad expressing Green mKikGR-CETN2 was stained with PlasMem Bright Green, followed by photoconversion of Green mKikGR-CETN2 to Red mKikGR-CETN2. 3D time-lapse images of the photoconverted germ cell were acquired every 10 minutes with z-sections at 1.5 μm intervals. 3D reconstructions were generated at 5-hour intervals over a total of 30 hours and are displayed in perspective. Cell surface 3D images are shown in green with overlays of reconstructed images at 10-80 minutes intervals. The trajectory of photoconverted Red mKikGR-CETN2 signals is represented using a color bar indicating time-lapse. See also Figure S10B and Movie 11. (G) Comprehensive time-lapse counts of photoconverted mKikGR-CETN2 foci during oocyte formation. Green mKikGR-CETN2 in a germ cell within cultured gonads/ovaries was photoconverted to Red mKikGR-CETN2, followed by 3D time-lapse imaging at 10-minute intervals. The number of Red mKikGR- CETN2 foci in photoconverted germ cells and adjacent non-photoconverted germ cells was counted and aligned by developmental age. *Ex vivo* time points were converted to corresponding *in vivo* developmental ages, as shown at the top of the plot. *N* = 13 photoconverted germ cells and 13 adjacent non-photoconverted germ cells. See also Figure S11. (H) Summary of changes in photoconverted mKikGR-CETN2 foci in adjacent non-photoconverted germ cells. The number of Red mKikGR-CETN2 foci, denoted as x, in each non-photoconverted germ cell adjacent to 13 photoconverted germ cells (G) was classified into two categories: *x* = 0 (no Red mKikGR-CETN2 foci detected throughout the observation) and *x* > 0 (at least one Red mKikGR-CETN2 focus detected at any time point during the observation). *N* = 13 adjacent non-photoconverted germ cells. Numbers in brackets indicate the sample sizes for each category. See also Figure S11.

### Absence of Detectable Centrosome Transfer Revealed by Photoconvertible Reporter Imaging

For this purpose, we generated mKikGR-CETN2 mice, in which CETN2 is tagged with the photoconvertible monomeric Kikume Green-Red (mKikGR) reporter (Figure S10A) (Habuchi et al., 2008). Immunostaining confirmed strong colocalization of mKikGR-CETN2 with endogenous CETN2 in germ cells (94.0%), validating its utility for centrosome recognition (Figure 5C-D). Exposure to 405 nm light successfully converted mKikGR fluorescence from green to red, enabling specific labeling of centrins within targeted germ cells (Figure 5E). Live imaging of photoconverted Red mKikGR-CETN2 in single germ cells allowed continuous tracking of centrin movement in 3D over 30 hours. While centrins exhibited dynamic movement within individual germ cells, no intercellular transfer to adjacent, non-photoconverted germ cells was observed (Figure 5F; Figure S10B; Movie 11). Additionally, the number of photoconverted Red mKikGR-CETN2 foci occasionally fluctuated within single cells, with some foci splitting into two to four or merging into clusters (Figure S10C). While the biological significance of these events remains unclear, they likely contribute to the changes in centrosome numbers observed across stages. Comprehensive quantification further confirmed the absence of centrosome transfer (Figure 5G-H; Figure S11). Across multiple time points, totaling 199 hours of observation, including 13 photoconverted germ cells and 13 adjacent non-photoconverted germ cells, photoconverted Red mKikGR-CETN2 signals remained restricted to the initially targeted germ cells. No sustained increase in red fluorescence was detected in adjacent non-photoconverted germ cells, indicating that centrosomes, like mitochondria, do not undergo detectable intercellular transfer during oocyte formation. Our live imaging analyses using mitochondrial and centrosome markers provide no evidence supporting organelle transfer between germ cells, contradicting the model proposed by Lei and Spradling (Lei & Spradling, 2016). Instead, our findings align with recent preprint data reported by Zhang and colleagues, which demonstrated no detectable cytoplasmic transfer between a total of 100 cell membrane borders using an independent imaging approach (Zhang et al., 2025).

In summary, our findings highlight several key aspects of germ cell development in the *ex vivo* culture system (Figure 6). First, we confirmed that the system reliably recapitulates meiotic progression, evidenced by the formation of chromosome axes, as well as early folliculogenesis, resulting in the production of GV oocytes. Our live imaging analysis provided valuable insights into germ cell behavior, including spherical transformation, autonomous size expansion of individual germ cells, and synchronized cyst degeneration. Notably, we observed dynamic blebbing activity, particularly during the mitosis-to-meiosis transition, and demonstrated its regulation by meiotic initiation signals and ROCK activity. Furthermore, our investigation into organelle transfer during oocyte formation found no evidence for detectable mitochondrial or centrosome transfer between germ cells, challenging the previously proposed model of extensive cytoplasmic transfer during cyst breakdown (see Discussion). Together, these results underscore the utility of our *ex vivo* culture system in studying these processes in real time and offer a comprehensive understanding of female germ cell development.

**Figure 6.**
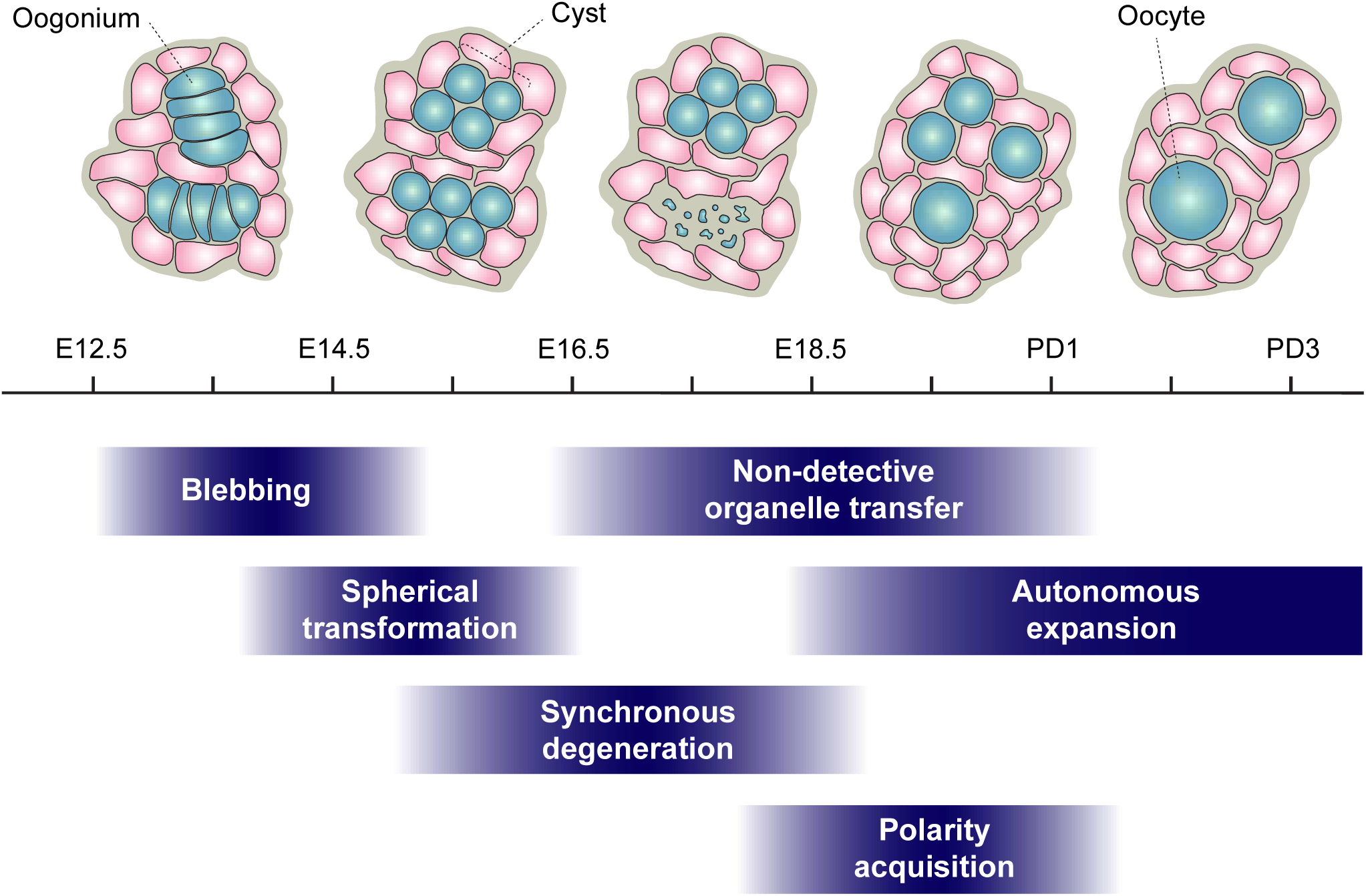
Summary of dynamic processes during oocyte formation. During the mitosis-to-meiosis transition, flattened oogonia exhibit prominent membrane blebbing, with the highest frequency observed at E14.5. As development progresses, germ cells undergo spherical transformation and form interconnected cysts. Synchronous germ cell degeneration was commonly observed within cysts, and separation of germ cells was also observed in other cysts. Between E16.5 and PD1, no detectable transfer of mitochondria or centrosomes was observed. During this period, germ cells acquire polarity and undergo substantial autonomous growth. Cyan, germ cells; Pink, somatic cells.

## Discussion

### Synchronous Germ Cell Degeneration and the Role of Intercellular Bridges

Our study revealed a strikingly synchronous pattern of germ cell degeneration during cyst breakdown in the fetal mouse ovary. This observation contrasts with the prevailing model of progressive, asynchronous attrition of nurse cells described in previous studies (Ikami et al., 2023; Lei & Spradling, 2016; Niu & Spradling, 2022). The collective degeneration, observed prominently between E15.5 and E18.5 in our live imaging analyses, suggests the existence of a coordinated mechanism that regulates cyst disassembly as a unit rather than through individual cell fate decisions.

We presume that this synchronized elimination is facilitated by the intercellular bridges that physically connect germ cells within each cyst (Ruby et al., 1969). Recent studies have highlighted regulatory functions of these structures in both male and female mouse germline development. In the male germline, intercellular bridges are essential for coordinating meiotic progression and repressing transposable elements, as their absence leads to defects in DNA replication, synapsis, and increased genomic instability (Sorkin et al., 2025). In females, intercellular bridges have been shown to coordinate the transition from mitosis to meiosis through dilution of regulatory factors across the cyst (Soygur et al., 2021), implying that they contribute to the synchronization of developmental transitions. More recently, Vasilev and colleagues demonstrated that long-lived cytokinetic bridges in early mouse embryos mediate the simultaneous elimination of sister cells (Vasilev et al., 2025). Their study showed that pro-apoptotic proteins such as Bax and caspase-3 were able to diffuse into the directly connected sister cell, but not into non-sister cells. These reports support the idea that intercellular bridges in germline cysts may facilitate the coordinated activation of cell death pathways or structural collapse, resulting in the near-simultaneous degeneration of multiple cyst cells.

These observations prompt a broader comparative question: Why do *Drosophila* germline cysts, which are also connected by intercellular bridges, show asynchronous cyst cell degeneration? One possible explanation lies in fundamental differences in cyst architecture and cytoskeletal organization between species. In *Drosophila*, intercellular bridges are transformed into large, actin-rich ring canals that connect to the fusome, a specialized organelle that establishes polarity, regulates cell cycles, and directs material transfer toward the oocyte (Hinnant et al., 2020). This organization enables selective cytoplasmic transport and may constrain the diffusion of apoptotic signals, thereby supporting the stepwise removal of nurse cells. In contrast, mouse germ cell cysts lack a canonical fusome and appear to rely more heavily on direct cytoplasmic coupling and shared signaling through bridges. This structural difference may permit broader and more uniform propagation of death signals, leading to the synchronous degeneration we observed.

Importantly, our data does not exclude the previously reported asynchronous mode of germ cell degeneration (Ikami et al., 2023; Lei & Spradling, 2016; Niu & Spradling, 2022). Rather, we propose that synchronous elimination represents an additional and possibly underrecognized mode of cyst breakdown during mouse oocyte formation. The coexistence of both synchronous and asynchronous degeneration may reflect distinct developmental contexts, intrinsic asymmetries among cysts, or temporally regulated shifts in how germ cell fates are resolved. Further investigation will be necessary to determine how these distinct modes are integrated and regulated during ovarian development.

### Rethinking Organelle Transfer and Cytoplasmic Enrichment

A prevailing model in mouse germ cell development posits that organelles are transferred from nurse cells to the prospective oocyte during cyst breakdown (Spradling et al., 2022). These organelles include Golgi complexes, mitochondria, and centrosomes, and the transfer is thought to occur between cyst cells connected by intercellular bridges. This model, largely extrapolated from studies in *Drosophila* and supported primarily by static observations in mouse ovaries, suggests that cytoplasmic and organelle transfer underlies the enrichment of cellular components critical for oocyte growth (Lei & Spradling, 2016; Niu & Spradling, 2022). However, our live imaging–based analysis challenges this view.

Using photoconvertible fluorescent reporters for mitochondria and centrosomes, we examined organelle dynamics within germ cell clusters during cyst breakdown. Although we observed active intracellular organelle movement, we did not detect directional transfer of either mitochondria or centrin-labeled centrosomes between neighboring germ cells. These findings suggest that, within the developmental window examined (E16.5 to E19.5/PD0) and under the spatial and temporal resolution of our imaging system, intercellular organelle transfer is unlikely to play a major role in cytoplasmic enrichment. While we cannot exclude the possibility of rare or subtle transfer events below the detection threshold, the robust detection of photoconverted signals within targeted cells argues against extensive transfer. Also, it remains possible that other classes of molecular cargo, such as RNAs or proteins, are selectively exchanged among germ cells without accompanying bulk organelle movement.

In *Drosophila*, intercellular bridges expand to form ring canals, which facilitate cytoplasmic transfer toward the oocyte (Robinson & Cooley, 1996). In contrast, in mice, the diameter of intercellular bridges reportedly decreases between E14.5 and E19.5/PD0 (Lei & Spradling, 2016). Lei and Spradling proposed the existence of plasma membrane gaps—structurally distinct from intercellular bridges—that might mediate organelle exchange between germ cells. However, the mechanism underlying the formation of these membrane gaps, as well as their presence and dynamics during cyst breakdown, remains poorly characterized. These insights highlight fundamental structural differences between *Drosophila* and mouse germ cell cysts, as well as potentially divergent mechanisms of organelle transfer during oocyte development.

A recent preprint proposed a distinct mechanism of cytoplasmic enrichment in which dominant oocytes engulf remnants of degenerated germ cells through an autophagy-related process (Zhang et al., 2025). While our study was not designed to directly monitor such engulfment events, this model aligns more closely with our observation that organelles remain confined to individual cells during cyst breakdown. It also leaves open the possibility that organelle transfer may occur via engulfment following cyst dissociation. In summary, our findings suggest that active organelle transfer between germ cells is unlikely to be the primary mechanism for cytoplasmic enrichment. Instead, the absence of detectable transfer highlights the need to revisit the models of oocyte formation and consider alternative pathways for cytoplasmic enrichment in oocytes. By enabling real-time observation of germ cell behavior, the imaging system established in this study uncovers previously inaccessible aspects and opens new directions for investigating the mechanisms underlying mammalian oocyte development.

## Materials and Methods

### Mice

Mouse lines used in this study were obtained from the following sources: C57BL/6NCrSlc (Japan SLC, Inc.); Stella-ECFP (accession no. CDB0465T, RIKEN) (Ohinata et al., 2008); H2B-mCherry (also referred to as R26- H2B-mCherry; accession no. CDB0239K, RIKEN) (Abe et al., 2011); mCherry-SYCP3 (Erasmus MC) (Enguita-Marruedo et al., 2018); Golgi-EGFP (also referred to as R26-Golgi-EGFP; accession no. CDB0246K, RIKEN) (Abe et al., 2011); Mito-EGFP (also referred to as R26-Mito-EGFP; accession no. CDB0251K, RIKEN) (Abe et al., 2011); PhAM^floxed^ (strain no. 018385, JAX) (Pham et al., 2012); EGFP-CETN2 (strain no. 008234, JAX) (Higginbotham et al., 2004); Spo11-Cre (strain no. 032646, JAX) (Lyndaker et al., 2013). The mKikGR-CETN2 mouse line (also referred to as R26-mKikGR-CETN2; accession no. CDB0470E, RIKEN) was generated in this study and is referred to hereinafter. Mito-Dendra2 mice were generated by crossing PhAM^floxed^ mice with Spo11-Cre mice to excise the stop sequences flanked by loxP sites. mCherry-SYCP3 mice, originally on an FVB background, were backcrossed to C57BL/6NCrSlc mice. Timed-pregnant females were obtained by natural mating of transgenic males aged 8 to 24 weeks with either wild-type C57BL/6NCrSlc females or transgenic females aged 7 to 16 weeks. The presence of a copulatory plug was designated as 0.5 days post coitum. All animal experiments were conducted in accordance with the ethical approvals of the Institutional Animal Care and Use Committee at RIKEN Kobe Branch.

### Ex vivo Culture

Gonads or ovaries were harvested from female fetuses between E11.5 and E15.5, with the mesonephros separated using 30-gauge needles. Fetal sex was determined by examining the gonad or ovary under a stereo microscope or by PCR genotyping of the fetal mesonephros, targeting the *Sry* and *Xist* genes, as previously described (Aizawa et al., 2020). For *ex vivo* culture, a 4-well cover glass chamber (5222-004, IWAKI) or a Lab-Tek chambered cover glass (155383, Thermo Fisher Scientific) was coated with Cell-Tak (354240, Corning) according to the manufacturer’s protocol. Each well of the chamber was overlaid with 190 µl of culture medium for the IWAKI chamber or 170 µl for the Lab-Tek chamber. The culture medium consisted of αMEM (21444-05, Nacalai Tesque) supplemented with 10% FBS (A3161002, Thermo Fisher Scientific), 10 mM HEPES (15630080, Thermo Fisher Scientific), 55 µM beta-mercaptoethanol (21985023, Thermo Fisher Scientific), and 50 U/ml penicillin-streptomycin (15070063, Thermo Fisher Scientific). Each gonad or ovary was placed at the center of a well and carefully transferred to an incubator maintained at 37°C with 5% CO₂ in air. After 12-18 hours, the medium was replaced with 200-240 µl of fresh culture medium to ensure the gonad or ovary remained slightly submerged. The culture was sustained by refreshing the medium every other day until further experimentation.

As appropriate for each experiment, the cell membrane, nucleus, or filamentous actin was stained using PlasMem Bright Green/Red (P504/P505, Dojindo Laboratories), Hoechst 33342 (H3570, Thermo Fisher Scientific), or SiR-Actin (CY-SC001, Cytoskeleton Inc), respectively. The staining reagents were added to the culture medium approximately 8 hours before imaging at a 2000-fold dilution for PlasMem Bright Green/Red, at a final concentration of 1 µg/ml for Hoechst 33342 or at a final concentration of 0.5 µM for SiR-Actin. Before imaging, the medium volume in each well was adjusted to over 300 µl as needed. For live imaging sessions exceeding 4 hours, the surface of the medium was covered with mineral oil to minimize evaporation.

### Microscopy and Imaging

Imaging was performed using Zeiss LSM780, LSM880, and LSM900 confocal microscopes, all equipped with a GaAsP detector, a 40× C-Apochromat 1.2 NA water immersion objective lens (Zeiss), and an environmental chamber for CO₂ and temperature control. The LSM780 and LSM880 were operated with Zen (Black Edition), and the LSM900 was operated with Zen (Blue Edition). The LSM900 was used for Mito- Dendra2 imaging, while the LSM780 and LSM880 were used for all other imaging applications.

For long-term imaging of oocyte formation, a single z-plane of 512 × 512 pixel xy images, covering a field of 141.70 × 141.70 µm, was acquired at 10-minute intervals. For 3D characterization of germ and somatic cells, 31 confocal z-sections were acquired at 1-µm intervals, generating 512 × 512 pixel xy images that covered a total volume of 70.85 × 70.85 × 30.00 µm. For nuclear rotation imaging, a single z-plane of 512 × 512 pixel xy images, covering 26.57 × 26.57 µm, was acquired at 10-second intervals for at least 20 minutes. For blebbing imaging, a single z-plane of 512 × 512 pixel xy images, covering 53.14 × 53.14 µm, was acquired at 2- or 3-second intervals for at least 10 minutes. For Golgi-EGFP imaging, a single z-plane of 512 × 512 pixel xy images, covering 70.85 × 70.85 µm, was acquired at 4-minute intervals over a period of 72 hours. For Mito-EGFP imaging, a single z-plane of 512 × 512 pixel xy images, covering 53.14 × 53.14 µm, was acquired at 1-minute intervals for 14 hours. For Mito-Dendra2 imaging, photoconversion of the Dendra2 protein was performed using the Bleaching option in ZEN software. The cytoplasmic region of a single germ cell was selected, and photoconversion was induced by applying 50 iterations of irradiation with a 405 nm laser at 5% output. Following photoconversion, Mito-Dendra2 signals were tracked for 4D spatial and temporal imaging using an adapted version of a macro initially developed by the research group of Dr. Jan Ellenberg (Politi et al., 2018). Three confocal z-sections were acquired at 2-µm intervals, generating 512 × 512 pixel xy images that covered a total volume of 53.24 × 53.24 × 4.00 µm at 5-minute intervals. For EGFP-CETN2 imaging, 12 confocal z-sections were acquired at 1.5-µm intervals, generating 512 × 512 pixel xy images covering a total volume of 106.27 × 106.27 × 16.50 µm at 10-minute intervals over a 24-hour period. For mKikGR-CETN2 imaging, photoconversion of the mKikGR protein was performed using the Bleaching option in ZEN software. A centrin focus region within a single germ cell was selected, and photoconversion was induced by irradiation with a 405 nm laser at 1% output with 20 iterations. Following photoconversion, 13 confocal z-sections were acquired at 1.5-µm spacing, generating 512 × 512 pixel xy images covering a total volume of 106.27 × 106.27 × 18.00 µm at 10-minute intervals.

### Characterization of Germ and Somatic Cells

For three-dimensional analysis, confocal z-stack images were reconstructed using Imaris software (Bitplane). Cell and nuclear surfaces were segmented by manually delineating boundaries in each z-section based on PlasMem Bright Red and Hoechst signals, respectively. Parameters including cell volume, cell sphericity, and nuclear sphericity were measured from these segmentations. The distance between the nuclear center and the cell surface center was calculated based on their spatial coordinates derived from the 3D reconstructions.

For 2D analysis, imaging data were processed using Fiji software (https://fiji.sc/). Cell boundaries were manually delineated by tracing the PlasMem Bright Red signal. Cellular parameters including cell circularity, cell area, and Stella-ECFP intensity were measured based on these boundaries. Nuclear boundaries were defined using the H2B-mCherry signal, and nuclear circularity, nuclear area, and H2B-mCherry intensity were quantified accordingly. Measurements obtained from Fiji were exported and analyzed using GraphPad Prism (GraphPad Sofware). Quantitative data were plotted at each time point to assess changes in cellular parameters over the course of the culture period.

### Long-term Analysis of Germ Cell Growth and Degeneration

Cell boundaries of tracked germ cells were manually delineated by tracing the PlasMem Bright Green signal using Fiji software, and cell area was quantified. To evaluate area expansion trends, a linear approximation was fitted to each time series using GraphPad Prism, and the slope of the fitted line was used as an indicator of the expansion rate. Developmental time was defined as the number of hours elapsed since E0.5, which corresponds to noon on the day a copulatory plug was detected in the female and was treated as a continuous variable based on the actual imaging time.

To analyze germ cell degeneration, we identified degeneration events in germ cells expressing H2B- mCherry based on chromatin condensation or fragmentation. For each developmental window between E13.5 and PD1, 12 regions of 100 μm × 100 μm were randomly selected from six E12.5 gonads cultured *ex vivo*. All degeneration events within these areas were manually counted. Germ cells undergoing degeneration in immediate proximity were considered part of the same cluster, and the number of germ cells per cluster was defined as the cluster size. Cluster sizes were plotted as individual data points and visualized using box and whisker plots generated in GraphPad Prism. Also, degenerated germ cells were categorized based on cluster size into four groups (single cells, 2-cell clusters, 3-cell clusters, and >4-cell clusters), and the cumulative number of germ cells in each category was calculated for each developmental window.

### Nuclear Rotation Assay

To assess nuclear rotation, wild-type E12.5 gonads were cultured *ex vivo* for four days and subsequently stained with Hoechst 33342 to visualize chromatin. Time-lapse imaging was performed at 10-second intervals using the Zeiss LSM880 confocal microscope, as described above. In each germ cell nucleus, specific foci within Hoechst-dense regions, presumed to correspond to heterochromatin, were manually selected as reference points. The movement of these foci was tracked across all time points using Imaris software, and their displacement and velocity were quantified at each time point. The resulting values were plotted over time.

### Blebbing Assay

To assess membrane blebbing dynamics in germ cells, female gonads from Stella-ECFP embryos at E11.5– E12.5 were cultured *ex vivo* and incubated with PlasMem Bright Red (Dojindo Laboratories) at a 1:2000 dilution for approximately 8 hours prior to imaging. Where indicated, gonads were also stained with SiR-Actin (Cytoskeleton Inc.) at a final concentration of 0.5 μM to visualize filamentous actin. Live imaging was performed at 2- or 3-second intervals using LSM780 or LSM880 confocal microscopes, as described above. Blebs were defined as membrane protrusions exceeding 2 μm in height, and the number of blebs was manually counted from the live imaging data.

For temporal projection analysis, membrane outlines were traced across seven consecutive time points (200 s total) using Adobe Illustrator (Adobe Inc.). To investigate the role of actin in blebbing, the bleb arc length was measured during both the expansion and contraction phases. This measurement was performed using Fiji software by manually tracing the arc at 3-second intervals. The distance between the arc ends was measured, along with the mean intensity of PlasMem Bright Red and SiR-Actin, which were plotted over time.

Pharmacological reagents were added to the culture medium either at the onset of culture or 16 hours prior to imaging, depending on the experimental design. The reagents used included LDN193189 (500 nM), BMS493 (10 μM), Jasplakinolide (10 nM-1 μM), Cytochalasin D (31.3-500 nM), Ciliobrevin D (6.3-100 μM), CK666 (10-250 μM), SMIFH2 (10-250 μM), Blebbistatin (1-25 μM), and H1152 (12-300 nM). Following treatment, blebbing frequency was quantified as described above.

### Immunostaining and Optical Clearing

Whole-mount immunostaining and optical clearing of fetal gonads or ovaries were conducted based on the SeeDB2 protocol, with minor modifications (Ke et al., 2016). Tissues were fixed in 4% paraformaldehyde (PFA) in PBS for 2 hours at 4°C, followed by three washes with PBS. Permeabilization was performed overnight at 4°C in PBS containing 2% saponin. Samples were then incubated in blocking buffer (FP1012, PerkinElmer) for 3 hours at 4°C. Primary antibodies were diluted in the blocking buffer and incubated with the tissues overnight at 4°C. After three washes with washing buffer (PBS containing 2% saponin and 0.5% Triton X-100), secondary antibodies diluted in the blocking buffer were applied and incubated overnight at 4°C.

For tissue clearing, samples were sequentially incubated in a 1:2 mixture of Omnipaque 350 (GE HealthCare) and Milli-Q water containing 2% saponin (solution 1) for 4-6 hours, followed by a 1:1 mixture (solution 2) for 6-12 hours. Clearing was completed by incubation in 100% Omnipaque 350 with 2% saponin and DAPI (0.1 µg/ml) for counterstaining (solution 3), for at least 12 hours. Cleared tissues were mounted in solution 3 on glass-bottom dishes. Images were acquired using a Zeiss LSM880 confocal microscope and processed with ZEN lite software (Zeiss).

The primary antibodies and their dilutions were anti-CETN2 (15877-1-AP, Proteintech; 1:400), anti-MVH (ab27591, abcam; 1:400), anti-STRA8 (provided by Dr. Kei-ichiro Ishiguro; 1:1000) (Shimada et al., 2023), anti-SCP3 (ab15093, abcam; 1:400), and anti-TEX14 (18351-1-AP, Proteintech; 1:500). The secondary antibodies and their dilutions were Alexa Fluor 488 goat anti-mouse IgG (A11029, Thermo Fisher Scientific; 1:500), Alexa Fluor 555 goat anti-mouse IgG (A21424, Thermo Fisher Scientific; 1:500), Alexa Fluor 568 goat anti-rabbit IgG (A11036, Thermo Fisher Scientific; 1:500), Alexa Fluor 647 donkey anti-mouse IgG (A31571, Thermo Fisher Scientific; 1:500), and Alexa Fluor 647 donkey anti-rabbit IgG (A31573, Thermo Fisher Scientific; 1:500).

### Mitochondria Transfer Assessment

To evaluate mitochondrial transfer between germ cells, we used Mito-Dendra2 mice expressing a photoconvertible mitochondrial reporter. Fetal gonads or ovaries were cultured *ex vivo* and stained with PlasMem Bright Green to label membranes. A single germ cell was photoconverted using a 405-nm laser to shift Mito-Dendra2 fluorescence from green to red, followed by 3D time-lapse imaging at 5-minute intervals using a Zeiss LSM900 confocal microscope, as described above. Photoconverted cells were tracked using the red Mito-Dendra2 signal. For each time point, a single Z-plane showing the maximum area of the targeted cell was selected, and cell boundaries were manually delineated using Fiji software. Mean red fluorescence intensity was quantified within both the photoconverted cell and neighboring, non- photoconverted germ cells. These intensity values were plotted over developmental time. Developmental time was determined based on the time elapsed from E0.5, defined as noon on the day a copulatory plug was detected in the mother, and was treated as a continuous variable based on the actual time of observation. To assess trends in red signal intensity in adjacent cells, an approximate line was fitted to each time series using Graphpad Prism, and its slope (*m*) was evaluated as an indicator of potential mitochondrial transfer. Slopes were categorized into two groups: a positive slope (*m* ≥ 0), indicating a mitochondrial transfer, and a negative slope (*m* < 0), indicating no transfer.

### Fluorescence Recovery after Photobleaching (FRAP)

Gonads were collected from E12.5 fetuses generated by crossing C57BL/6NCrSlc mice with EGFP-CETN2 transgenic mice. After seven days of *ex vivo* culture (E12.5 + 7d), germ cells exhibiting EGFP-CETN2 signals were selected for FRAP analysis. Photobleaching was performed by exposing a defined region of EGFP- CETN2 signal to a 488 nm laser at 4% output for 15 seconds. A circular region of interest (ROI) with a diameter of 2 µm was placed over the bleached area to measure fluorescence recovery. The mean EGFP- CETN2 fluorescence intensity within the ROI was quantified every 5 minutes for a total of 120 minutes using Fiji software. Fluorescence intensity values were normalized to a baseline value of 1, corresponding to the mean pre-bleach intensity.

### Generation of mKikGR-CETN2 Mice

The conditional mKikGR-CETN2 mouse line (also referred to as R26R-mKikGR-CETN2; accession no. CDB0395E, RIKEN; https://large.riken.jp/distribution/reporter-mouse.html) was generated via CRISPR/Cas9-mediated knock-in in zygotes, following the method described by Abe and colleagues (see Figure S10A) (Abe et al., 2020). To construct the pCMV-mKikGR-CETN2 vector, the mKikGR cDNA sequence was excised from the mKikGR-C1 vector (Addgene #54656) and ligated into the hCent2-pEGFP-C1 vector (Addgene #29559) after digestion with *AgeI* and *BspEI* restriction enzymes. The donor vector contains 5’ (900 bp) and 3’ (600 bp) homology arms targeting intron 1 of the ROSA26 locus, an adenovirus splicing acceptor (SA), a stop cassette flanked by loxP sites, and a bovine growth hormone polyadenylation (bpA) sequence (Srinivas et al., 2001). The mKikGR-CETN2 cDNA fragment was excised from pCMV-mKikGR- CETN2 and inserted between the stop cassette and bpA sequences to complete the donor vector. The guide RNA (gRNA) was designed to target a site downstream of exon 1 in the ROSA26 locus (5’- CGCCCATCTTCTAGAAAGAC-3’). A mixture of Cas9 protein (100 ng/μl) (Thermo Fisher Scientific), crRNA (50 ng/μl), tracrRNA (100 ng/μl) (FASMAC), and the donor vector (10 ng/μl) was microinjected into the pronuclei of C57BL/6N one-cell zygotes. After microinjection, zygotes were transferred into the oviduct of 0.5 dpc pseudopregnant females for implantation. A total of 28 F0 founder mice were obtained, of which 21 (8 females and 13 males) were identified as correctly targeted based on PCR genotyping. The following primers were used for PCR analysis:

Wild-type allele (217 bp): P1 (5’-TCCCTCGTGATCTGCAACTCCAGTC-3’) and P2 (5’- AACCCCAGATGACTACCTATCCTCC-3’) R26R allele (270 bp): P1 (5’-TCCCTCGTGATCTGCAACTCCAGTC-3’) and P3 (5’-GCTGCAGGTCGAGGGACC-3’) mKikGR insertion (543 bp): P4 (5’-ATCGAGCTGCGTATGGAAGG-3’) and P5 (5’-GTCTGGCAACTGGACAACCT- 3’)

To generate the constitutive mKikGR-CETN2 mouse line (also referred to as R26-mKikGR-CETN2; accession no. CDB0470E, RIKEN), two males carrying the targeted sequence were selected and crossed with Spo11- Cre female mice. Genotyping PCR was performed on F1 mice using the following primers, designed to amplify the R26 allele (606 bp): P6 (5’-GCTCAGTTGGGCTGTTTTGG-3’) and P7 (5’- CCTTCCATACGCAGCTCGAT-3’). A total of 10 F1 mice (6 females and 4 males) were confirmed to have the R26 allele based on PCR genotyping and sequencing. Among these, two males were selected for breeding to establish the mKikGR-CETN2 mouse colony.

### Centrosome Transfer Assessment

To assess centrosome transfer between germ cells, we utilized mKikGR-CETN2 mice expressing a photoconvertible centrin reporter. Fetal gonads or ovaries were cultured *ex vivo* and stained with PlasMem Bright Green to visualize cell membranes. A centrin signal within a single germ cell was photoconverted from green to red using a 405-nm laser on Zeiss LSM780 or LSM880 confocal microscopes, as described above. Three-dimensional time-lapse imaging was performed at 10-minute intervals with 1.5 μm z-sections for up to 30 hours. For 3D reconstruction, time-lapse image stacks of the photoconverted germ cell were processed using Imaris software. The cell border was manually delineated in each z-section based on the PlasMem Bright Green signal. Three-dimensional reconstructions of the cell surface were generated at selected time intervals between 10 and 80 minutes. Red mKikGR-CETN2 signals were detected at every time point and visualized using the Dragon Tail function in Imaris to represent the temporal trajectory of photoconverted centrosomes. To evaluate potential centrosome transfer, Red mKikGR-CETN2 foci were manually counted in both the photoconverted germ cell and a neighboring, non-photoconverted germ cell at each time point using Imaris software and aligned to developmental time. Developmental time was determined based on the time elapsed from E0.5, defined as noon on the day a copulatory plug was detected in the mother, and was treated as a continuous variable based on the actual time of observation.

### Statistical Analysis

Statistical analyses were performed using GraphPad Prism 9. A two-tailed, unpaired Welch’s t-test was used to assess differences between groups. Sample sizes (*n*) and *P* values are provided in the corresponding figures and figure legends. Error bars in figures indicate mean ± standard deviation (SD). No statistical methods were used to predetermine sample sizes. The threshold for statistical significance was set at *P* < 0.05.

## Acknowledgements

We are grateful to Dr. Willy Baarends (Erasmus MC) for providing the mCherry-SYCP3 mouse line and to Dr. Mitinori Saitou (Kyoto University) for the Stella-ECFP mouse line. We thank the Kobe Imaging and Information Analysis Facility (KbiIF) at RIKEN BDR for technical support with imaging, and Ms. Shiori Ori (RIKEN BDR) for assistance with 3D image analysis. We are grateful to Dr. Ryuki Shimada and Dr. Kei-ichiro Ishiguro (Kumamoto University) for providing the STRA8 antibody and for their insightful comments and suggestions. We also acknowledge Dr. Katsuhiko Hayashi (Osaka University) and Dr. Genshiro Sunagawa (RIKEN BDR) for valuable discussions and suggestions. This work was supported by RIKEN Pioneering Project “Long-Timescale Molecular Chronobiology”, JSPS KAKENHI Grant Number 23H04948, 21H02407, 25H00981, and the Naito Foundation Research Grant.

---

**Figure S1.**
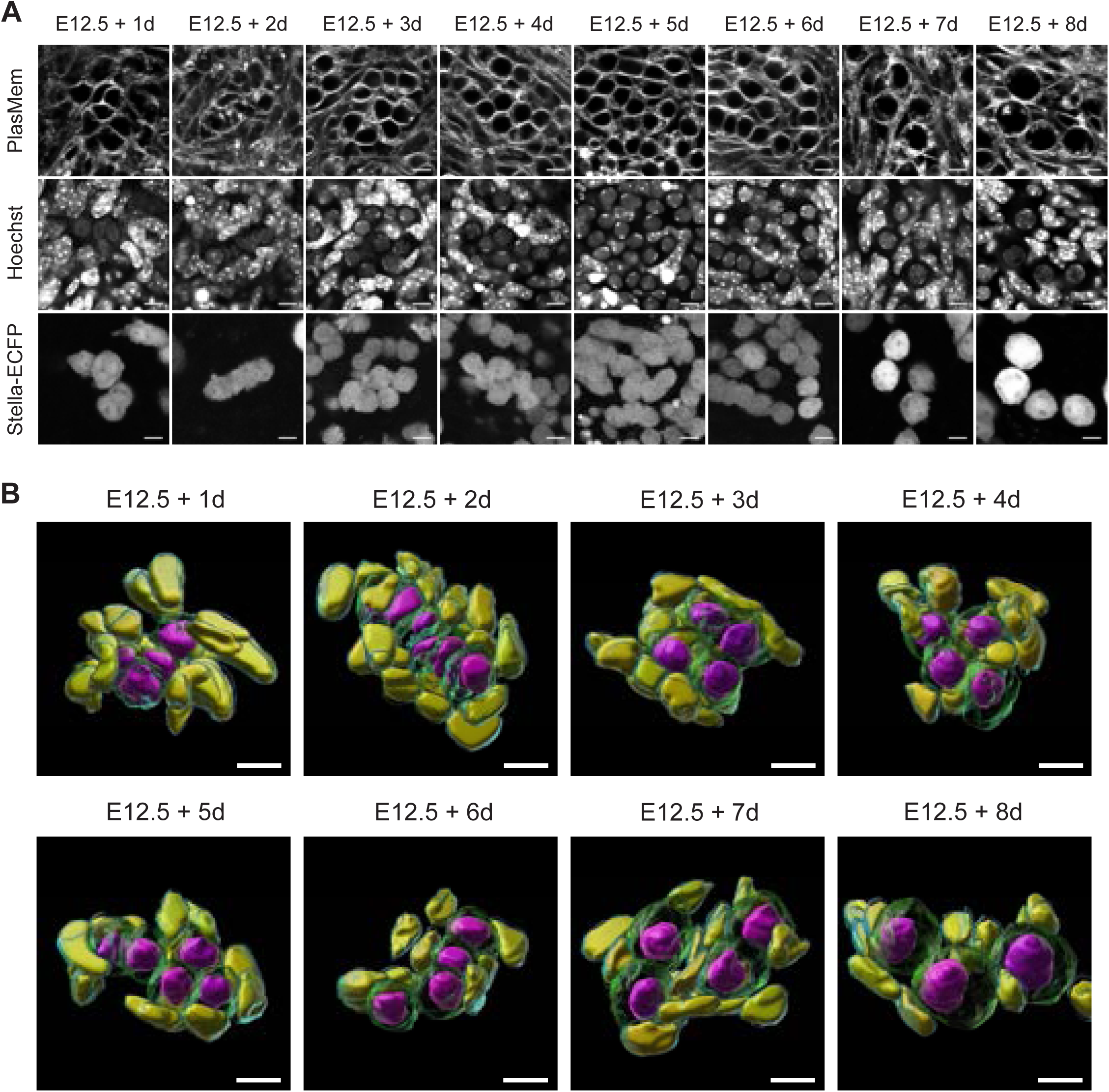
Three-dimensional reconstruction of germ and somatic cells in ex vivo culture. (A) Representative images of cells in E12.5 gonads expressing Stella-ECFP during an 8-day ex vivo culture. Gonads were stained with Hoechst 33342 and PlasMem Bright Red prior to imaging. Confocal z-sections were acquired at 1 µm intervals, and representative 2D sections are shown. Scale bar, 10 µm. (B) Three-dimensional reconstructed images of cells in E12.5 gonads expressing Stella-ECFP during an 8-day ex vivo culture. Gonads were stained with Hoechst 33342 and PlasMem Bright Red. Germ cell membranes (green), germ cell nuclei (magenta), somatic cell membranes (cyan), and somatic cell nuclei (yellow) are shown. Confocal z-sections were acquired at 1 µm intervals, and 3D images were reconstructed orthogonally. Identical images of E12.5 + 2d, E12.5 + 5d, and E12.5 + 8d gonads are shown in Figure 1D. Scale bar, 10 µm.

**Figure S2.**
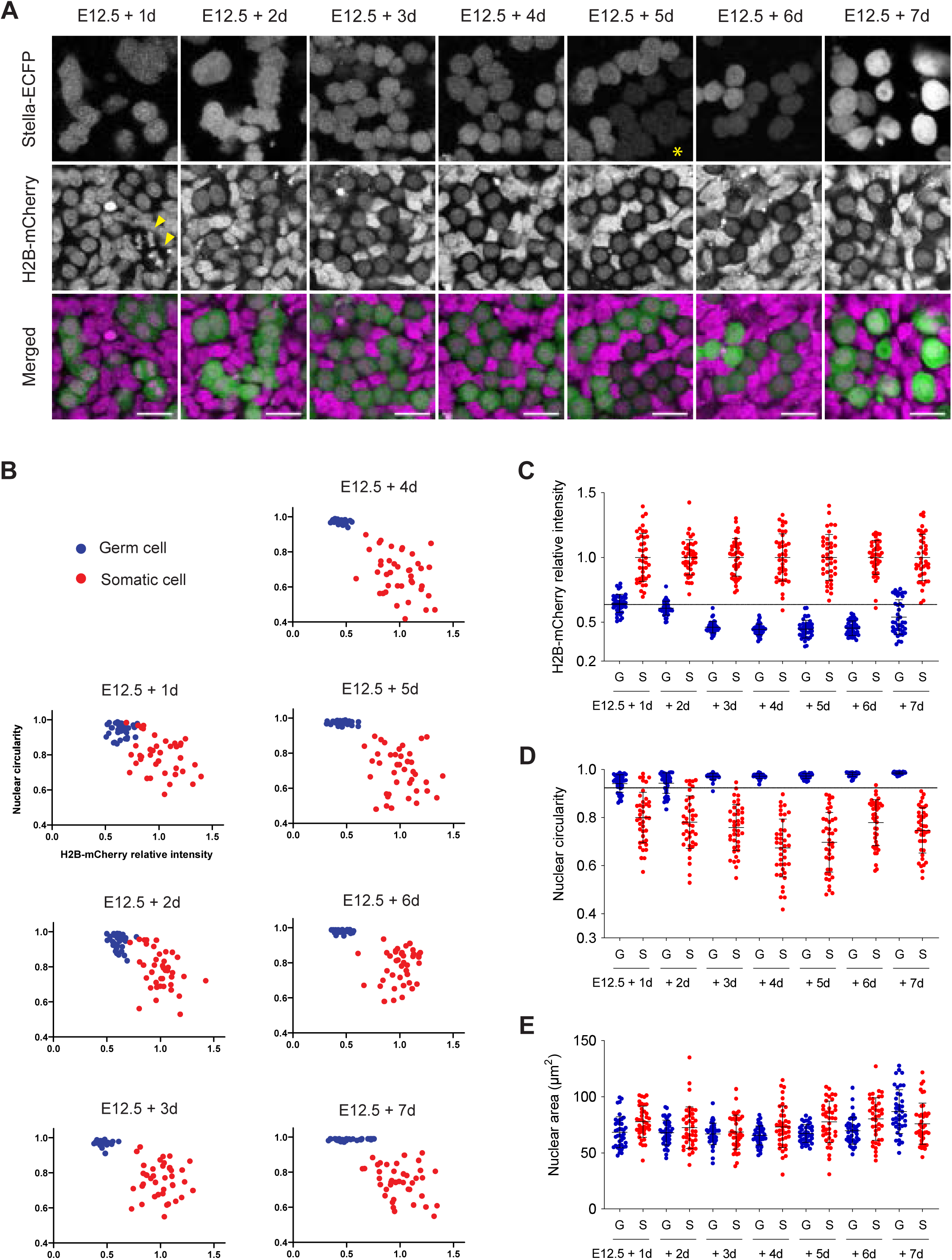
Characterization of germ and somatic cells in ex vivo culture. (A) Development of germ cells in E12.5 gonads co-expressing Stella-ECFP and H2B-mCherry during a 7-day ex *vivo* culture. The merged image (bottom) shows Stella-ECFP (green) and H2B-mCherry (magenta) expression. Stella-ECFP signals weakened around day 5 of culture but strengthened by day 7. An asterisk indicates cells with weakened Stella-ECFP signals. Arrowheads mark Stella-ECFP-positive cells undergoing presumed anaphase chromosome segregation. Scale bar, 20 µm. (B) Quantitative analysis of histone intensity and nuclear circularity in cells from E12.5 gonads co-expressing Stella-ECFP and H2B-mCherry during the 7-day *ex vivo* culture. From E12.5 + 3d to E12.5 + 7d, two distinct populations were identified: germ cells with low H2B-mCherry intensity and high nuclear circularity, and somatic cells with high H2B-mCherry intensity and low nuclear circularity. The mean H2B-mCherry intensity of somatic cells was normalized to a relative intensity of 1. (C, D, E) Quantitative analysis of histone intensity (C), nuclear circularity (D) and nuclear area (E) in cells from E12.5 gonads co-expressing Stella-ECFP and H2B-mCherry during the 7-day *ex vivo* culture. Bars represent mean values ± standard deviations. (C) The mean H2B-mCherry intensity of somatic cells was normalized to a relative intensity of 1. A dashed line indicates a H2B-mCherry relative intensity threshold of 0.63, distinguishing somatic cells (>0.63; 98.8% (158/160)) and germ cells (≤0.63; 100% (160/160)) between E12.5 + 3d and E12.5 + 6d. (D) A dashed line indicates a nuclear circularity threshold of 0.93, distinguishing somatic cells (>0.93; 99.0% (198/200)) and germ cells (≤0.93; 99.5% (199/200)) between E12.5 + 3d and E12.5 + 7d. (B, C, D, E) Germ cells (blue) and somatic cells (red) were distinguished based on Stella-ECFP expression intensity. Quantification included 40 germ cells and 40 somatic cells at each developmental time point. G, germ cell; S, somatic cell.

**Figure S3.**
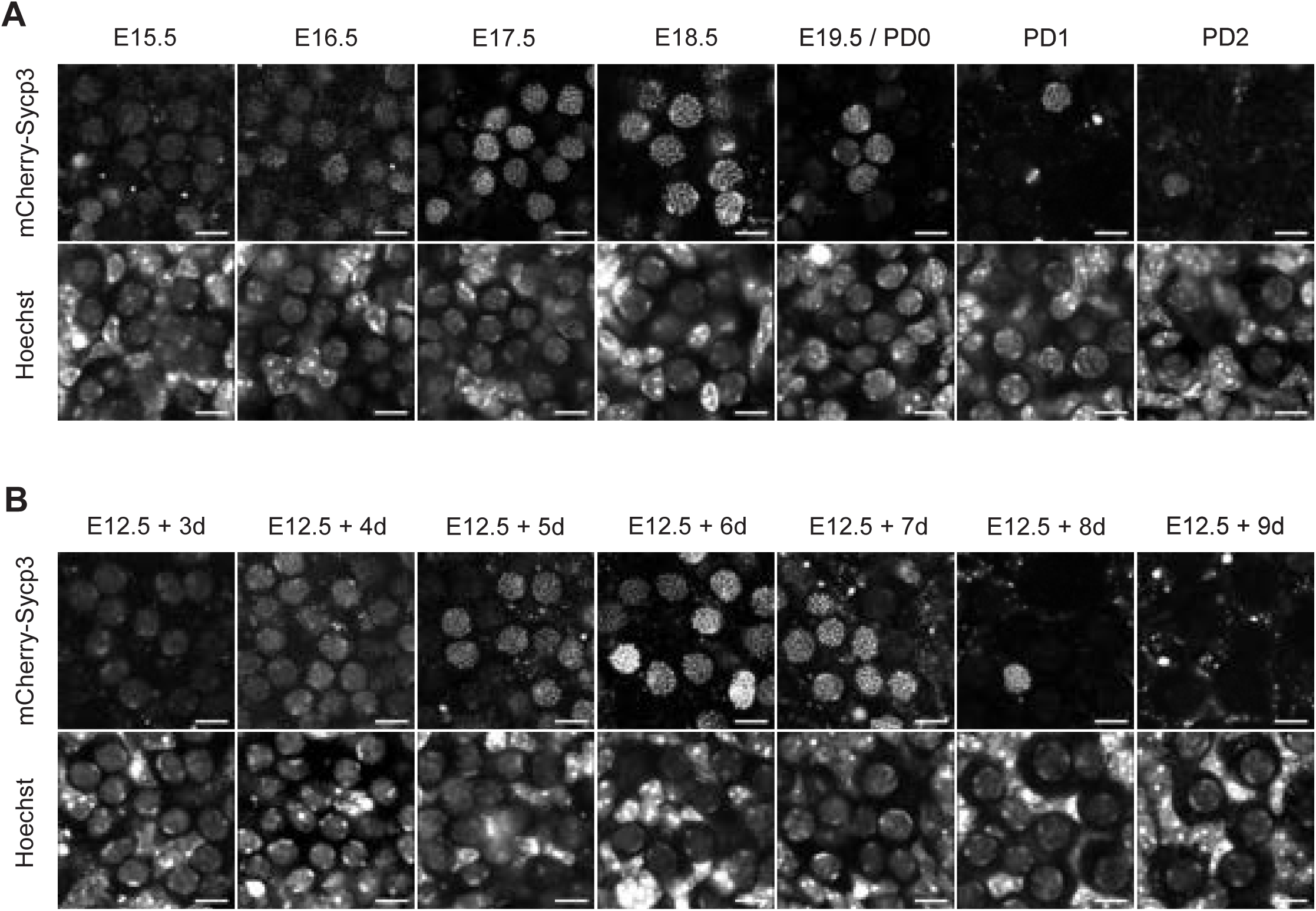
Comparison of chromosome axis formation in germ cells during in vivo and ex vivo development. (A) Representative images of germ cells in ovaries collected from E15.5 fetuses to PD2 neonates. Ovaries expressing mCherry-SYCP3 were stained with Hoechst 33342 prior to imaging. Chromosome axis formation was prominently observed between E17.5 and E19.5/PD0. Identical images of an E18.5 ovary are shown in Figure 2A. Scale bar, 10 µm. (B) Representative images of germ cells in E12.5 gonads during a 9-day ex *vivo* culture. E12.5 gonads expressing mCherry-SYCP3 were stained with Hoechst 33342 prior to imaging. Chromosome axis formation was prominently observed between E12.5 + 5d and E12.5 + 7d. Identical images of an E12.5 + 6d gonad are shown in Figure 2A. Scale bar, 10 µm.

**Figure S4.**
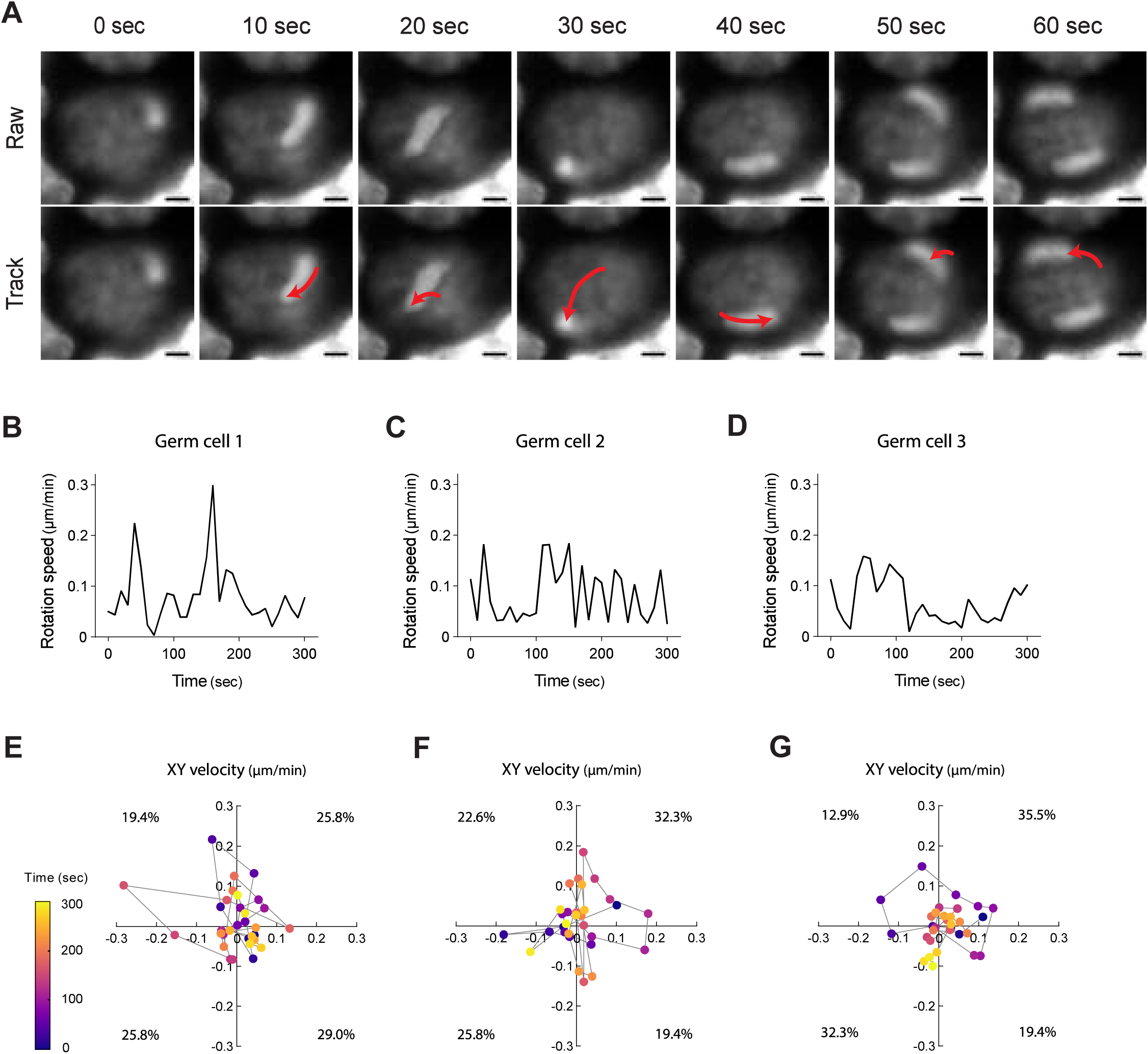
Characterization of nuclear rotation dynamics in germ cells. (A) Representative live imaging of a germ cell nucleus in an E12.5 + 4d gonad cultured ex vivo. The gonad was stained with Hoechst 33342 and imaged at 10-second intervals. Raw and tracking images are shown, with red lines indicating trajectories of nuclear rotation. Scale bar, 2 µm. See also Movie 2. (B-D) Time-lapse quantification of nuclear rotation speed for three representative germ cells. Rotation speed was measured every 10 seconds over a 300-second period. (E-G) Plots of X velocity (x-axis) versus Y velocity (y-axis) recorded at 10-second intervals for a 300-second sequence. Each plot corresponds to the germ cells analyzed in (B), (C), and (D), respectively. The number within each quadrant indicates the proportion of XY velocity plots among the 30 measurements.

**Figure S5.**
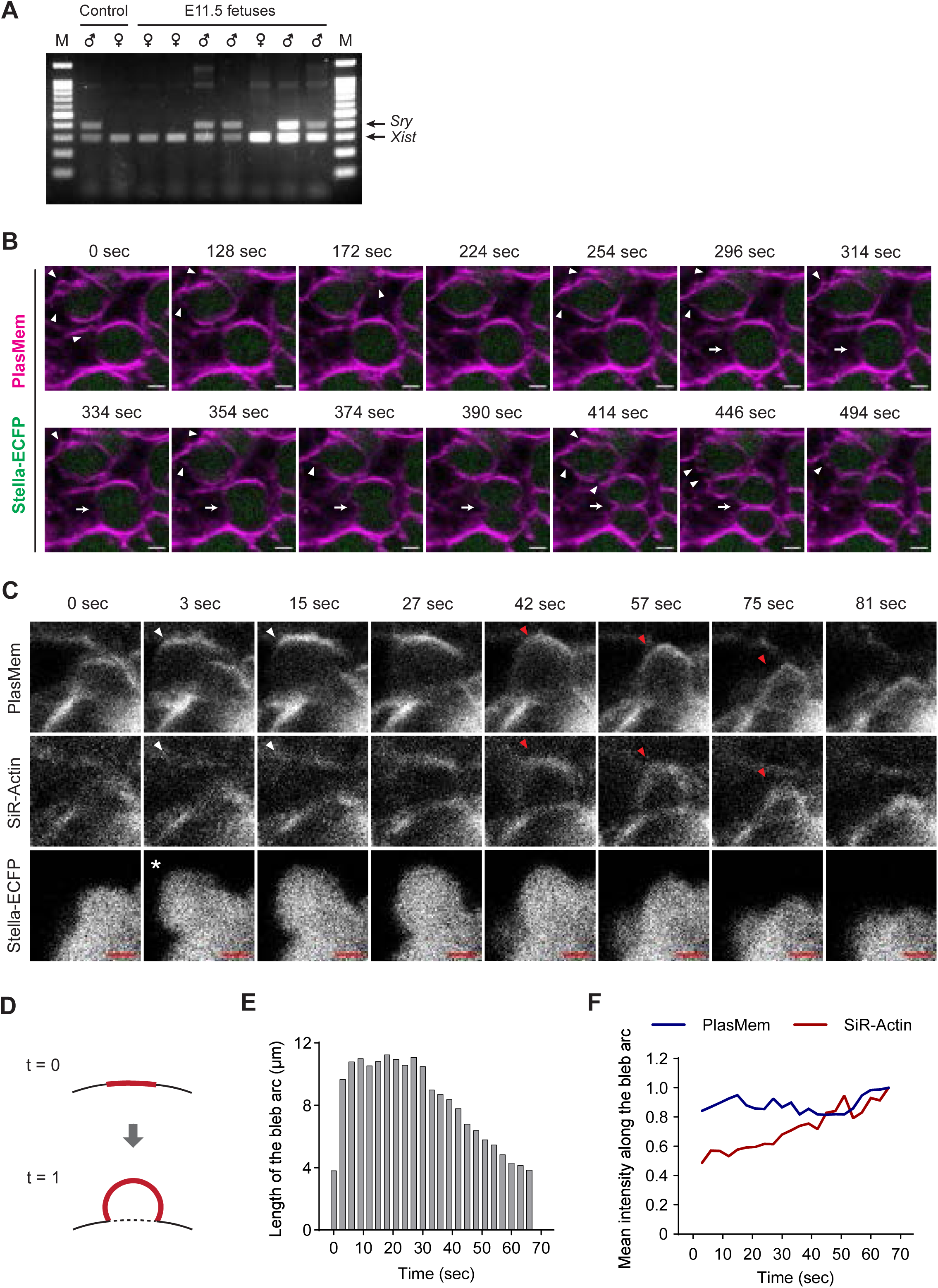
Characterization of blebbing in germ cells. (A) PCR analysis of E11.5 fetuses using Sry and Xist primers to determine fetal sex. M, marker. (B) Live imaging of germ cells in an E12.5 gonad expressing Stella-ECFP after 2 days of ex vivo culture. The gonad was stained with PlasMem Bright Red and imaged every 2 seconds. Stella-ECFP (green) and PlasMem (red) signals are shown as merged images. Arrowheads mark blebs, and arrows indicate the position of cytokinesis. Scale bar, 5 µm. See also Movie 4. (C) Representative live-imaging of blebbing in an E12.5 gonad expressing Stella-ECFP after 1 day of ex vivo culture. The gonad was stained with PlasMem Bright Red and SiR-Actin. Images were captured every 3 seconds. An asterisk marks an emerging bleb. White arrowheads indicate cell membranes with weak SiR-Actin signals, while red arrowheads denote membranes with distinct SiR-Actin signals. Scale bar, 2 µm. See also Movie 5. (D-F) Quantitative analysis of the blebbing shown in (C). (D) Schematic illustration of the measurement. The red line indicates the bleb arc, representing the measured linear region of interest. (E) Quantification of the bleb arc length over time. The arc length decreases after approximately 30 seconds. (F) Mean intensities of PlasMem Bright Red (blue) and SiR-Actin (red) along the bleb arc over time. PlasMem intensity remained stable, while SiR-Actin intensity increased as the bleb shrank.

**Figure S6.**
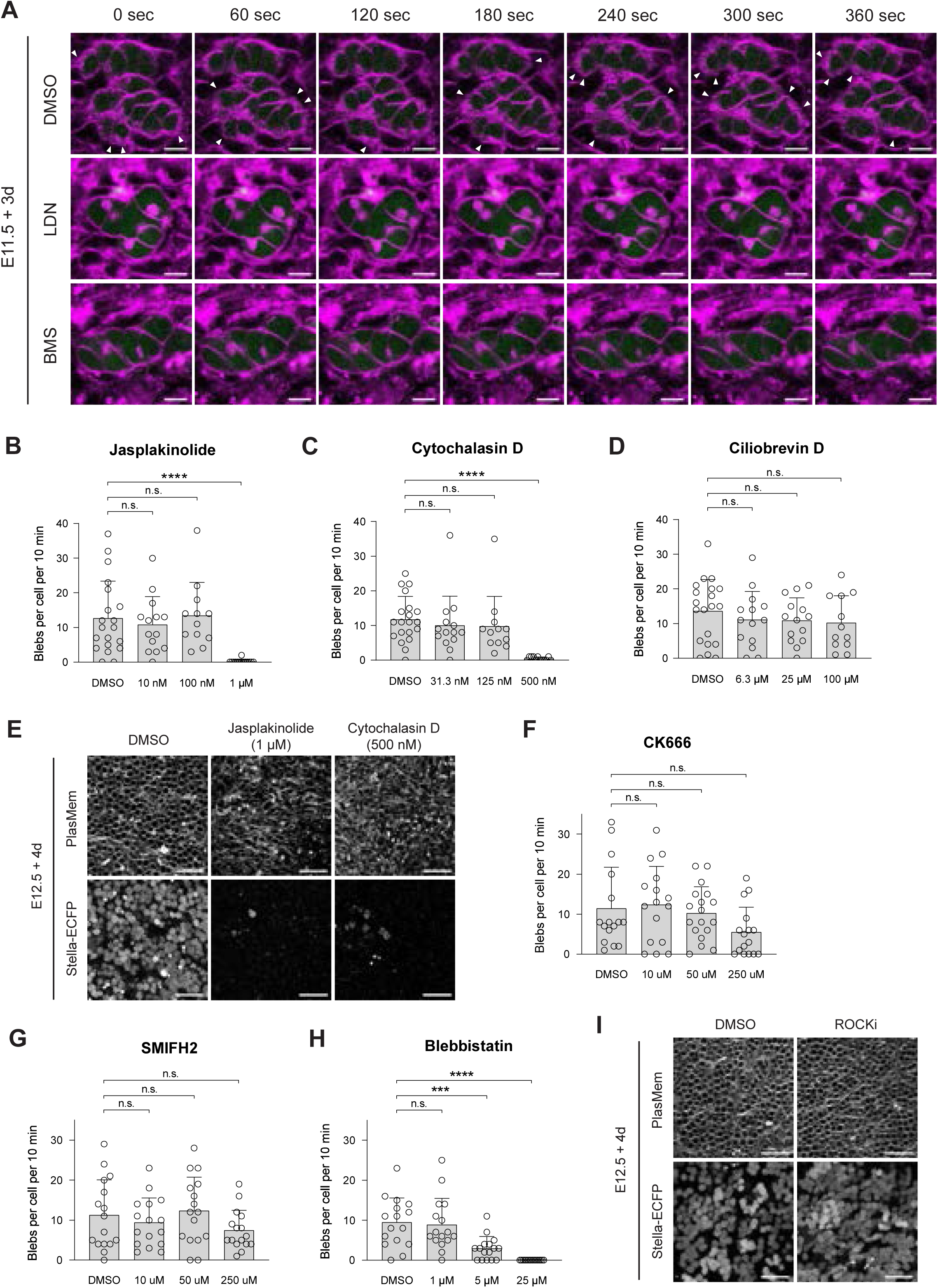
Analysis of signaling pathways regulating blebbing in germ cells. (A) Representative live imaging of E11.5 + 3d gonads under inhibition of meiotic initiation signals. Female E11.5 gonads were cultured for 3 consecutive days with DMSO, LDN1931189 (500 nM) or BMS493 (10 µM) treatments, followed by staining with PlasMem Bright Red. Arrowheads indicate blebs. Scale bar, 10 µm. See also Movie 6. (B-D) Blebbing frequency following treatment with Jasplakinolide (B), Cytochalasin D (C), or Ciliobrevin D (D). The number of blebs was counted from live imaging of E12.5 + 2d gonads expressing Stella-ECFP. Each inhibitor was supplemented 16 hours prior to imaging. Bars represent mean values + standard deviations. Statistical analysis was performed using a t-test with Welch’s correction. Sample sizes: (B) DMSO (n = 20), 10 nM (n = 14), 100 nM (n = 13), 1 µM (n = 14); (C) DMSO (n = 20), 31.3 nM (n = 14), 125 nM (n = 12), 500 nM (n = 12); (D) DMSO (n = 20), 6.3 µM (n = 14), 25 µM (n = 14), 100 µM (n = 12). **** P < 0.0001; ns, non-significant. (E) Representative images of cells in E12.5 + 4d gonads expressing Stella-ECFP. Gonads were cultured with DMSO, Jasplakinolide (1 µM), or Cytochalasin D (500 nM) for 16 hours between E12.5 + 1d and E12.5 + 2d, followed by staining with PlasMem Bright Red. Stella-ECFP-positive cells were abundant in samples treated with DMSO but were scarcely observed following treatment with Jasplakinolide or Cytochalasin D. Scale bar, 50 µm. (F-H) Blebbing frequency following treatment CK666 (F), SMIFH2 (G), or Blebbistatin (H). The number of blebs was counted from live imaging of E12.5 + 2d gonads expressing Stella-ECFP. Each inhibitor was supplemented 16 hours prior to imaging. Bars represent mean values + standard deviations. Statistical analysis was performed using a t-test with Welch’s correction. Sample sizes: (F) DMSO (n = 16), 10 µM (n = 16), 50 µM (n = 18), 250 µM (n = 16); (G) DMSO (n = 16), 10 µM (n = 16), 50 µM (n = 16), 250 µM (n = 16); (H) DMSO (n = 16), 1 µM (n = 16), 5 µM (n = 16), 25 µM (n = 16). *** P < 0.001; **** P < 0.0001; ns, non-significant. (I) Representative images of cells in E12.5 + 4d gonads expressing Stella-ECFP. Gonads were cultured with either DMSO or ROCKi (12 nM) for 16 hours between E12.5 + 1d and E12.5 + 2d, followed by staining with PlasMem Bright Red. Scale bar, 50 µm

**Figure S7.**
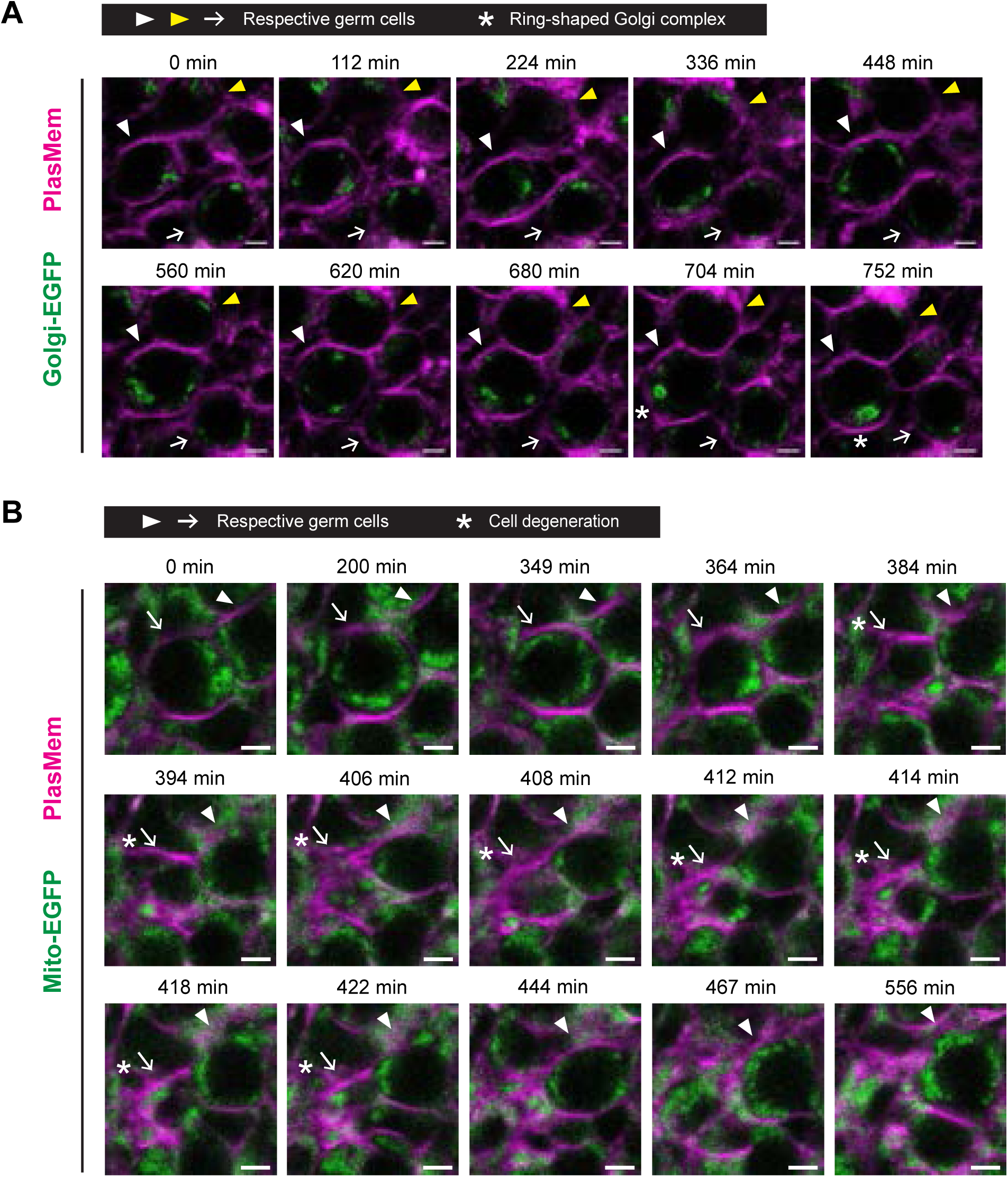
Live imaging of Golgi and mitochondrial dynamics during oocyte formation. (A) Representative live imaging of an E15.5 + 2d ovary expressing Golgi-EGFP. The ovary was stained with PlasMem Bright Red and imaged at 4-minute intervals. Golgi-EGFP (green) and PlasMem (magenta) signals are shown as merged images. Tracked germ cells are indicated by arrows, white arrowheads, and yellow arrowheads, respectively. Asterisks mark ring-shaped Golgi complexes, characteristic of the Balbiani body. Scale bar, 10 µm. See also Movie 7. (B) Representative live imaging of an E14.5 + 6d ovary expressing Mito-EGFP. The ovary was stained with PlasMem Bright Red and imaged at 1-minute intervals. Merged images show Mito-EGFP (green) and PlasMem (magenta) signals. Arrows and arrowheads indicate a tracked germ cell and its adjacent germ cell, respectively. Asterisks denote the degeneration of the tracked germ cell. Scale bar, 5 µm. See also Movie 8.

**Figure S8.**
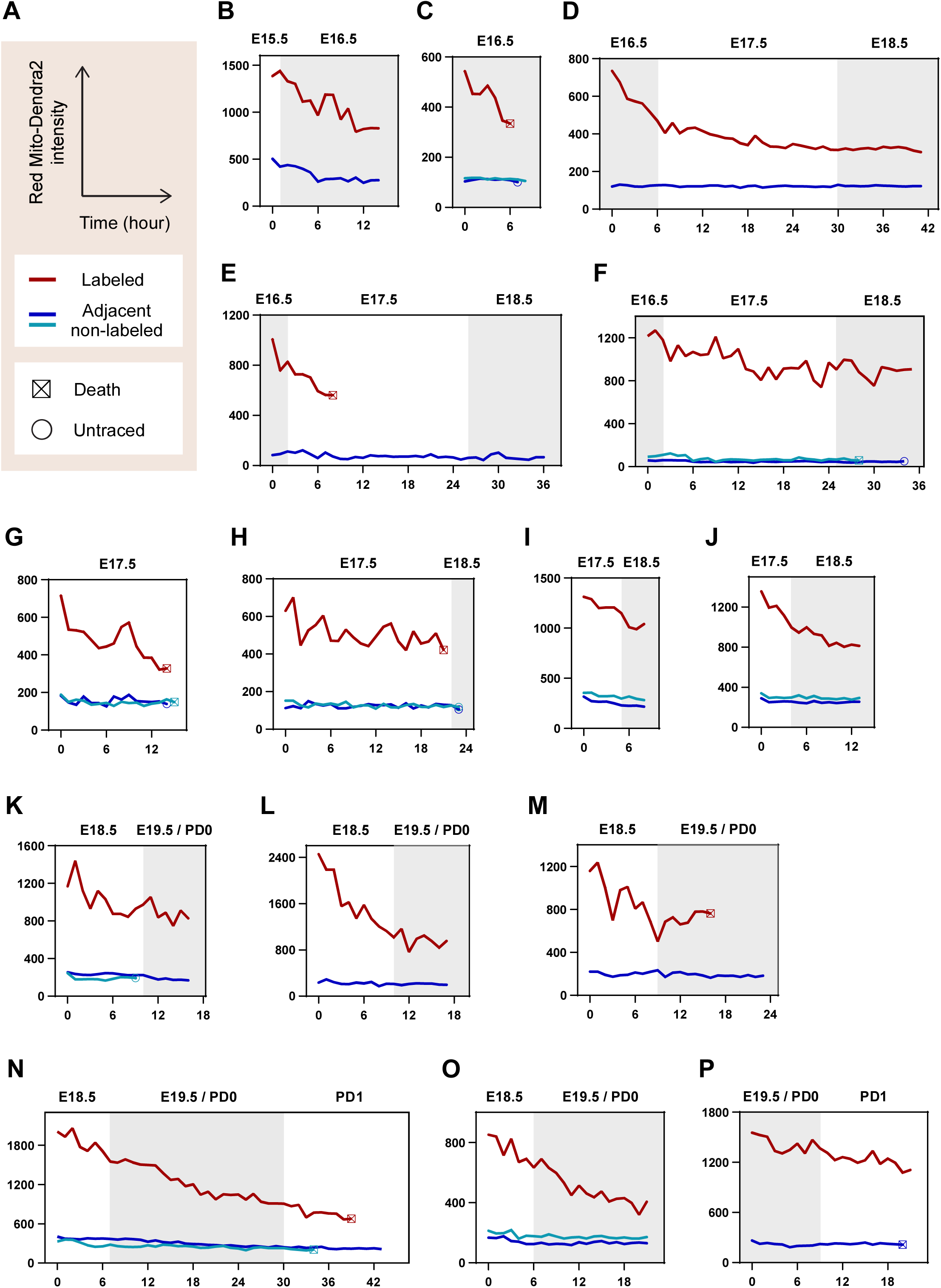
Individual time-lapse quantification of photoconverted Mito-Dendra2 intensity. Green Mito-Dendra2 in a germ cell within cultured gonads/ovaries was photoconverted to Red Mito-Dendra2, followed by 3D tracking and imaging at 5-minute intervals. Absolute mean intensities of Red Mito-Dendra2 in photoconverted germ cells, along with one or two adjacent non-photoconverted germ cells, were measured and aligned by developmental time. Ex vivo time points were converted to their corresponding in vivo developmental times, as shown at the top of each plot. (A) Axes titles and legends. (B-P) Plots of quantified Red Mito-Dendra2 intensity. The samples used for imaging include: (B) E12.5 + 3d gonad, (C) E12.5 + 4d gonad, (D) E12.5 + 4d gonad, (E) E14.5 + 2d ovary, (F) E14.5 + 2d ovary, (G) E12.5 + 5d gonad, (H) E12.5 + 5d gonad, (I) E12.5 + 5d gonad, (J) E12.5 + 5d gonad, (K) E12.5 + 6d gonad, (L) E12.5 + 6d gonad, (M) E12.5 + 6d gonad, (N) E12.5 + 6d gonad, (O) E12.5 + 6d gonad, (P) E12.5 + 7d gonad. See also Figure 4D-E.

**Figure S9.**
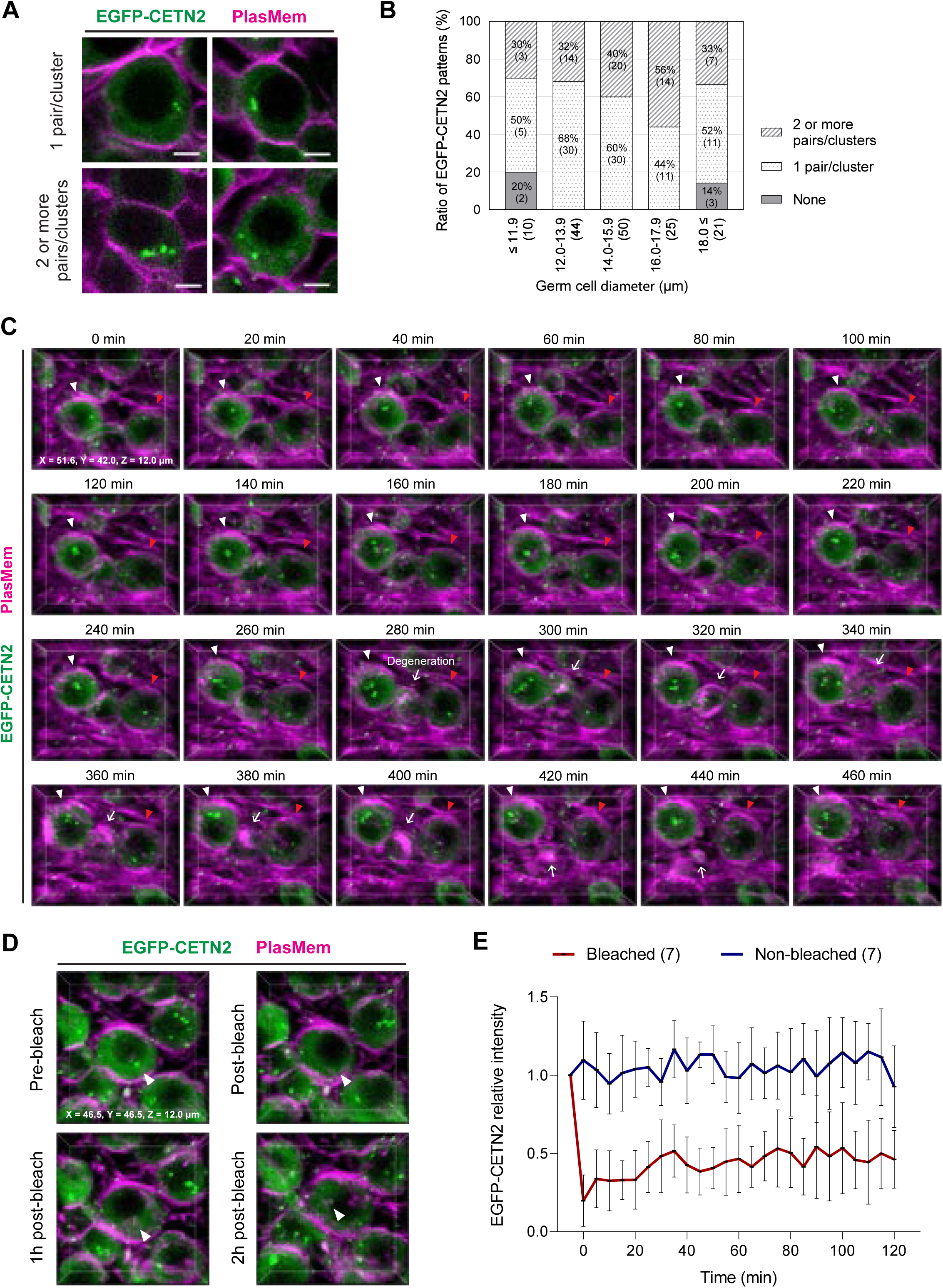
Characterization of EGFP-CETN2 dynamics during oocyte formation. (A) Patterns of EGFP-CETN2 signals in germ cells. Images were captured from E12.5 + 7d (bottom left) and E12.5 + 9d (top left, top right, and bottom right) gonads stained with PlasMem Bright Red. Merged images show EGFP-CETN2 (green) and anti-CETN2 (magenta). EGFP-CETN2 signal patterns include 1 pair (top left), 1 cluster (top right), and 2 or more pairs/clusters (bottom left and bottom right). Scale bar, 5 µm. (B) Distribution of EGFP-CETN2 patterns in germ cells by diameter. EGFP-CETN2 patterns in germ cells from E12.5 gonads cultured for 3 to 11 days were classified into three categories: None, 1 pair/cluster, and 2 or more pairs/clusters. Germ cell diameters were measured and grouped into five categories: ≤ 11.9 µm, 12.0 -13.9 µm, 14.0 -15.9 µm, 16.0 -17.9 µm, ≥ 18.0 µm. Numbers in brackets indicate germ cell counts. A total of 180 germ cells were analyzed, with 30 cells each from E12.5 + 3d, + 5d, + 7d, + 9d, and + 11d gonads. See also Figure 5A. (C) Representative live imaging of an E12.5 + 7d gonad expressing EGFP-CETN2. The gonad was stained with PlasMem Bright Red and subjected to 3D time-lapse imaging every 10 minutes with z-sections at 1.5 µm intervals. Merged 3D images show EGFP-CETN2 (green) and PlasMem (magenta) signals displayed every 20 minutes in perspective. White and red arrowheads indicate tracked germ cells, respectively, while arrows denote germ cell degeneration. See also Movie 10. (D, E) Assessment of EGFP-CETN2 photobleaching. E12.5 + 7d gonads expressing EGFP-CETN2 were stained with PlasMem Bright Red, followed by targeted photobleaching of EGFP-CETN2 signals in germ cells. (D) Representative 3D images of EGFP-CETN2 photobleaching. Merged images show EGFP-CETN2 (green) and PlasMem (magenta) signals, with arrowheads indicating the photobleached EGFP-CETN2 focus. (E) Time-lapse analysis of EGFP-CETN2 intensity in response to photobleaching. Mean EGFP-CETN2 intensity before photobleaching was normalized to a relative intensity of 1. The measurement area was manually defined using a circle with a diameter of 2 µm. The time point directly after photobleaching was set as 0 on the X-axis. Bars represent mean values ± standard deviations. N = 7 photobleached germ cells and 7 non-photobleached germ cells.

**Figure S10.**
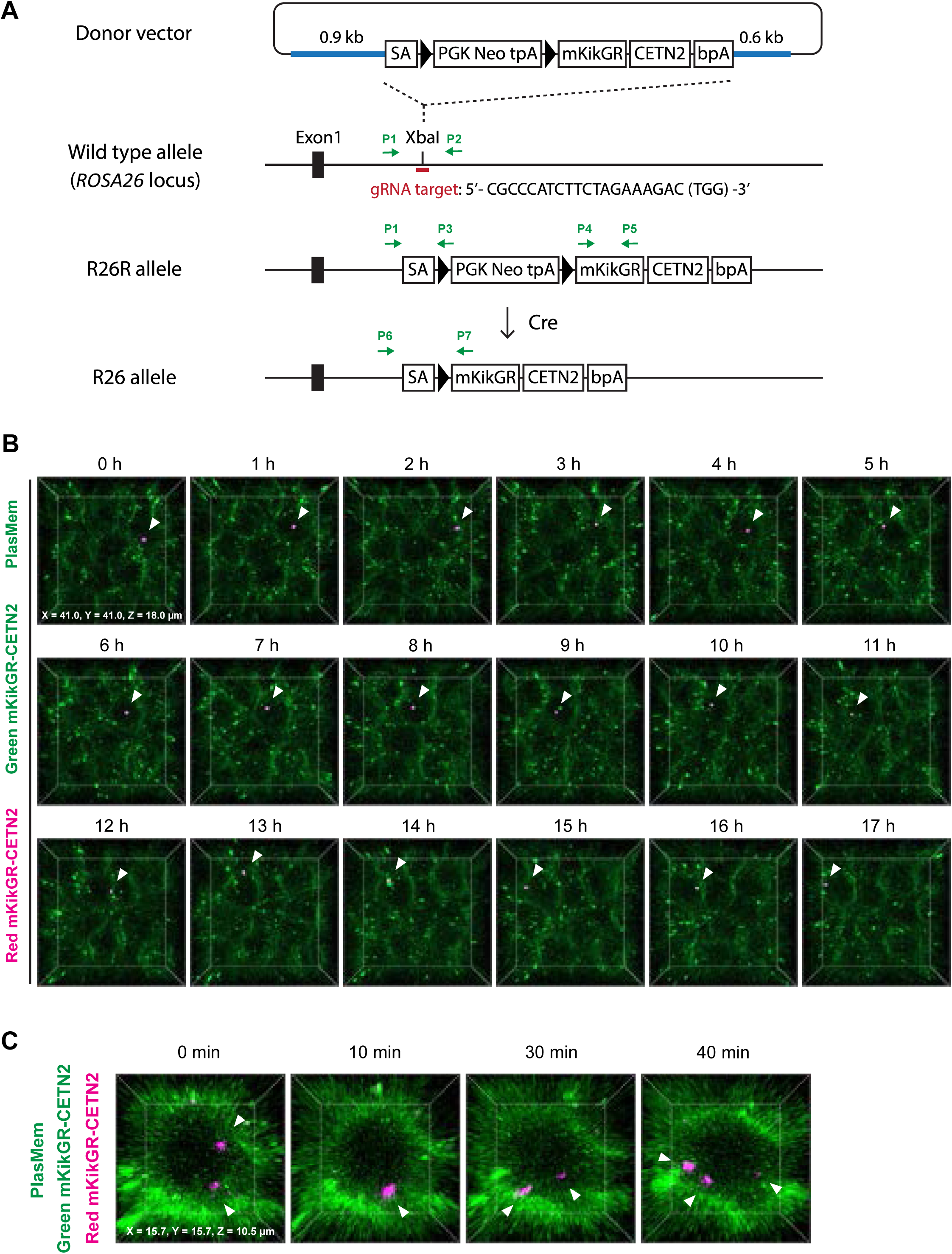
Generation of mKikGR-CETN2 mice and characterization of mKikGR-CETN2 signals. (A) Generation of mKikGR-CETN2 Mice. The donor vector contains 5’ (0.9 kb) and 3’ (0.6 kb) homology arms (blue) targeting intron 1 of the ROSA26 locus, flanking an expression cassette. The vector was inserted into the genome of C57BL/6N zygotes using the CRISPR-Cas9 system with a guide RNA targeting the ROSA26 genomic locus. The resulting mice were crossed with Spo11-Cre mice to excise the stop sequences flanked by loxP sites. The black box represents exon 1 of the Rosa26 gene, and black triangles indicate loxP sequences. Green arrows indicate PCR primers. Abbreviations: SA, adenovirus splice acceptor sequence; PGK Neo, neomycin resistance gene driven by the PGK1 promoter; tpA, triple repeats of the SV40 polyadenylation sequence; mKikGR, monomeric mKikGR Green-Red; bpA, bovine growth hormone polyadenylation sequence. (B) Representative time-lapse 3D Images of photoconverted Red mKikGR-CETN2 signals. An E12.5 + 5d gonad expressing Green mKikGR-CETN2 was stained with PlasMem Bright Green, followed by photoconversion of Green mKikGR-CETN2 to Red mKikGR-CETN2. 3D time-lapse images of the photoconverted and neighboring germ cells were captured every 10 minutes with z-sections at 1.5 µm intervals. In the merged images, PlasMem and non-photoconverted Green mKikGR-CETN2 are shown in green, while photoconverted Red mKikGR-CETN2 is shown in magenta. Arrowheads indicate photoconverted Red mKikGR-CETN2 signals. See also Figure 5F and Movie 11. (C) Representative images of changes in mKikGR-CETN2 focal count. An E12.5 + 5d gonad expressing mKikGR-CETN2 was stained with PlasMem Bright Green, followed by photoconversion of Green mKikGR-CETN2 to Red mKikGR-CETN2. 3D time-lapse images were captured every 10 minutes with z-sections at 1.5 µm intervals. Arrowheads indicate photoconverted Red mKikGR-CETN2, with focal counts varying over time. In merged images, PlasMem and non-photoconverted Green mKikGR-CETN2 are shown in green, while photoconverted Red mKikGR-CETN2 is shown in magenta.

**Figure S11.**
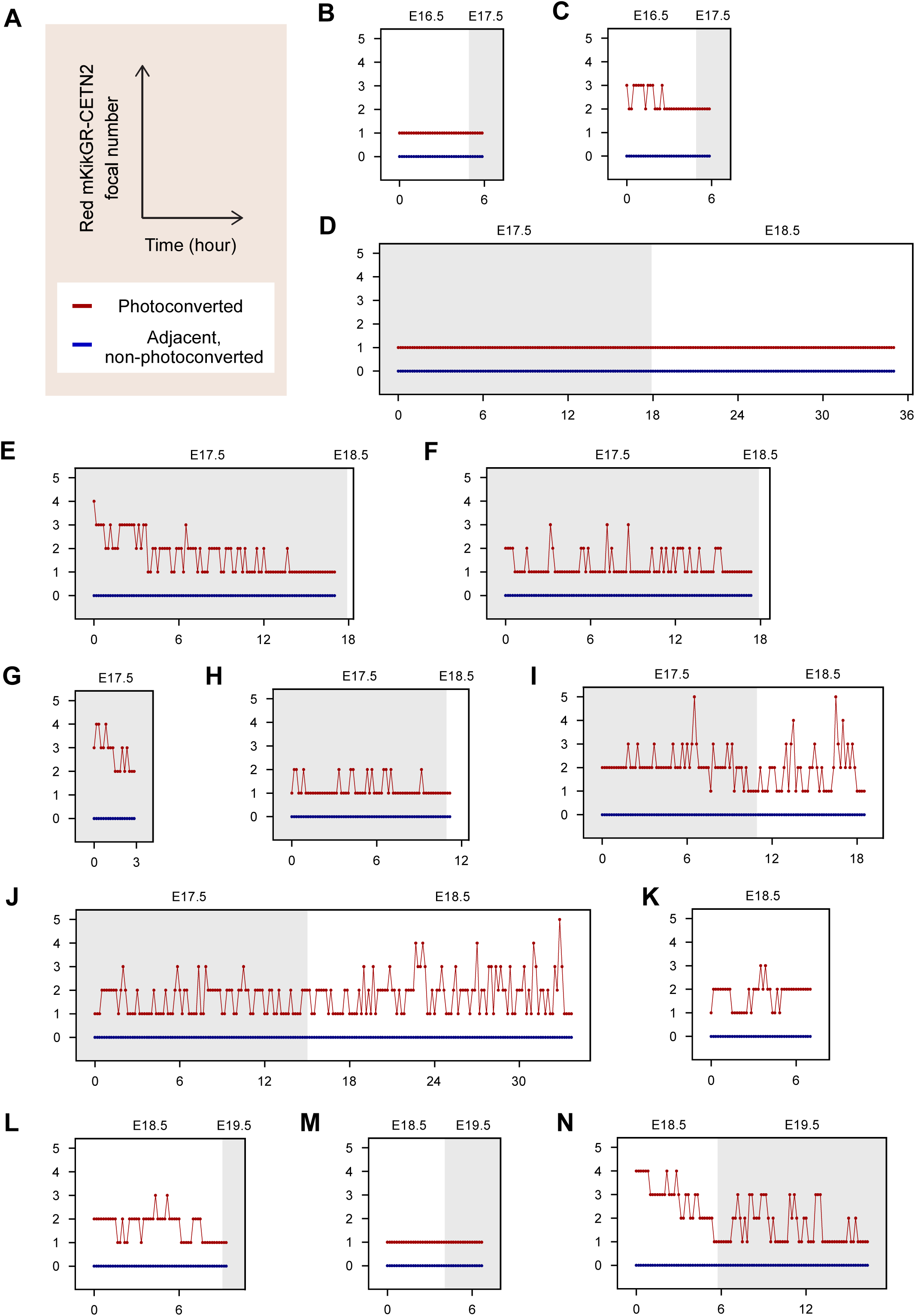
Individual time-lapse counts of photoconverted mKikGR-CETN2 foci. Green mKikGR-CETN2 in a germ cell within cultured gonads was photoconverted to Red mKikGR-CETN2, followed by 3D imaging at 10-minute intervals. The number of Red mKikGR-CETN2 foci in photoconverted germ cells and adjacent non-photoconverted germ cells was manually counted at each time point and aligned by developmental time. Ex vivo time points were converted to corresponding in vivo developmental times, as shown at the top of each plot. (A) Axes titles and legends. (B-N) Plots showing the number of Red mKikGR-CETN2 foci. The samples used for imaging include: (B) E12.5 + 4d gonad, (C) E12.5 + 4d gonad, (D) E12.5 + 5d gonad, (E) E12.5 + 5d gonad, (F) E12.5 + 5d gonad, (G) E12.5 + 5d gonad, (H) E12.5 + 5d gonad, (I) E12.5 + 5d gonad, (J) E12.5 + 5d gonad, (K) E12.5 + 6d gonad, (L) E12.5 + 6d gonad, (M) E12.5 + 6d gonad, and (N) E12.5 + 6d gonad. See also Figure 5G-H.

